# G Protein-Coupled Receptor-Ligand Dissociation Rates and Mechanisms from τRAMD Simulations

**DOI:** 10.1101/2021.06.20.449151

**Authors:** Daria B. Kokh, Rebecca C. Wade

## Abstract

There is a growing appreciation of the importance of drug-target binding kinetics for lead optimization. For G protein-coupled receptors (GPCRs), which mediate signaling over a wide range of timescales, the drug dissociation rate is often a better predictor of *in vivo* efficacy than binding affinity, although it is more challenging to compute. Here, we assess the ability of the τ-Random Acceleration Molecular Dynamics (τRAMD) approach to reproduce relative residence times and reveal dissociation mechanisms and the effects of allosteric modulation for two important membrane-embedded drug targets: the β2-adrenergic receptor and the muscarinic acetylcholine receptor M2. The dissociation mechanisms observed in the relatively short RAMD simulations (in which molecular dynamics (MD) simulations are performed using an additional force with an adaptively assigned random orientation applied to the ligand) are in general agreement with much more computationally intensive conventional MD and metadynamics simulations. Remarkably, although decreasing the magnitude of the random force generally reduces the number of egress routes observed, the ranking of ligands by dissociation rate is hardly affected and agrees well with experiment. The simulations also reproduce changes in residence time due to allosteric modulation and reveal associated changes in ligand dissociation pathways.

**TABLE OF CONTENTS GRAPHIC:** 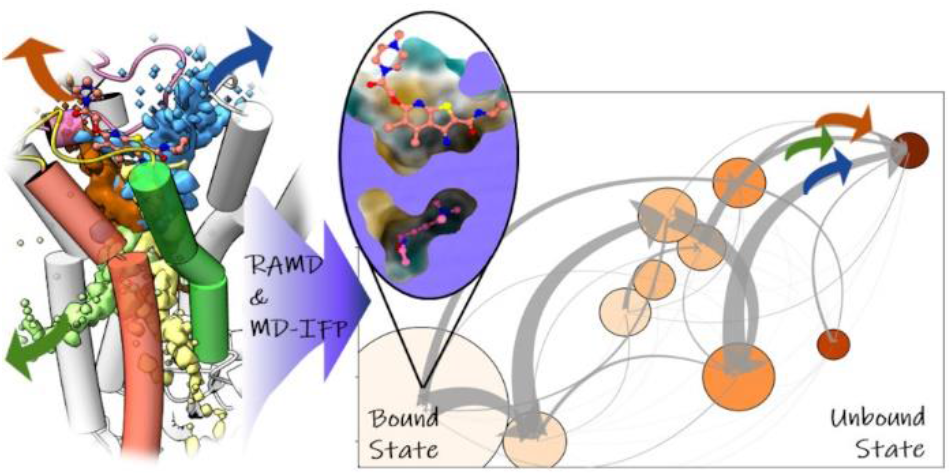

## 1. INTRODUCTION

G protein-coupled receptors, GPCRs, are targets for over a third of all clinical drugs, and these drugs are used for the treatment of a broad spectrum of diseases ^1^. The key function of GPCRs that is targeted by drugs is signal transduction through the cell membrane. Signal transduction is controlled by the binding of various orthosteric and allosteric compounds, ions and lipids to GPCRs, as well as by the ability of GPCRs to undergo structural transformations ^2^. The interplay between these dynamic processes regulates signal transmission in a time-dependent manner, indicating that particular signaling profiles may be achieved through the optimization of the GPCR binding kinetic parameters of drugs ^3,4,5,6,7^. The impact of the binding and unbinding kinetics on GPCR drug efficacy *in vivo* has been outlined in several recent reviews and is currently an area of active research^8,3,9,10^. Another important aspect of GPCRs is that all the members of this protein superfamily have a seven transmembrane α-helix bundle structure, which makes the design of drugs with high selectivity quite challenging^11^. Therefore, considerable effort has been devoted to discovering drugs that allosterically modulate GPCRs and can have a variety of effects on the behavior of orthosteric ligands, thus enabling discrimination between different downstream pathways and resulting in fewer side-effects. ^12,7,13,11^

Over the last few years, a large number of methods to compute binding kinetic rates have been published, many of which employ molecular dynamics (MD) simulation approaches ^14,15^. Despite a plethora of methods, the prediction of drug-protein dissociation rates (or their inverse: residence time, τ) by computation is still highly challenging and under methodological development. The main reasons for this are that simulation protocols often require extensive computational resources and high user expertise and that the results often lack accuracy. The long timescales of protein-drug dissociation events relative to conventional MD simulations mean that many methods make use of enhanced sampling MD-based approaches, which improve computational efficiency and make the calculation of unbinding rates for large numbers of compounds feasible. However, such methods may introduce biases in the simulation results and, therefore, their accuracy and their ability to reveal correct mechanistic insights into the dissociation process requires assessment.

We recently developed the τRAMD protocol for computing relative protein-ligand dissociation rates and exploring the mechanisms of protein-ligand unbinding ^16,17, 18^. The basis of τRAMD is the Random Acceleration MD (RAMD)^19^ simulation method, in which an additional randomly oriented force is applied in an adaptive manner to the ligand center of mass to facilitate the ligand unbinding process, thereby making simulation of dissociation of a drug-like compound possible on the nanosecond time-scale without *a-priori* knowledge of the egress route or mechanism or extensive parameter adjustment. The ability of the τRAMD method to give correct relative residence times of a large set of inhibitors was first evaluated and shown for inhibitors of Heat Shock Protein 90^16^ and it was subsequently shown to distinguish the kinetic selectivity of inhibitors of two closely related kinases ^20^. Recently, it was demonstrated that τRAMD not only enables computation of the relative τ values of small compounds unbinding from different cavities in several T4 lysozyme mutants at a range of temperatures but also reveals egress routes, and the metastable states along them, in very good agreement with those from conventional MD and other enhanced sampling MD studies^21^

In the present work, we assess the application of τRAMD to more complex systems, membrane-embedded GPCRs in their complexes with orthosteric and allosteric compounds, whose τ values span from seconds to minutes. Specifically, we study two experimentally well-characterized drug targets: the β2-adrenergic receptor (β2AR) and the muscarinic acetylcholine receptor M2 (mAChR M2). While RAMD has previously been applied to study ligand egress routes from GPCRs ^22,23^, the ability of τRAMD to predict relative residence times and identify the factors determining the residence times has not previously been assessed for membrane-embedded proteins with allosteric modulation.

The binding and unbinding of orthosteric β2AR ligands have been simulated in several computational studies. The unbinding of the small inverse agonist beta blocker, carazolol, from β2AR was first simulated using RAMD over a decade ago by Wang and Duan^23^. In this work, several egress channels were revealed, both between the transmembrane helices and through the extracellular vestibule, ECV. Subsequently, the binding of three antagonists (propranolol, alprenolol, and dihydroalprenolol) and one agonist (isoproterenol) to β2AR was explored in extensive conventional all-atom MD simulations^24^. Several spontaneous association events of the compounds to β2AR through the ECV were observed during a cumulative MD simulation time of over 100 µs. The free energy profile estimated was characterized by two energy barriers, corresponding to (1) ligand desolvation upon entrance into the ECV, and (2) ligand displacement into the orthosteric binding pocket. This study was further extended for dihydroalprenolol in Ref ^25^, where the advantages of adaptive sampling over traditional MD simulations in the identification of binding pathways were demonstrated. Finally, in Ref. ^26^, conventional MD simulations coupled with metadynamics simulations of alprenolol were employed to build a Markov State Model and find a Path Collective Variable. This enabled construction of a minimum free energy pathway of protein-ligand association which, although it did not reveal any transition barriers related to ligand desolvation, did show a shallow local energy minimum for the ligand positioned in the ECV.

mAChR M2, which is bound by the neurotransmitter agonist, acetylcholine (ACh), has long been a model system for studying allosteric regulation. The structures of the mAChR M2 complex with the Positron Emission Tomography superagonist, [^3^H]iperoxo (IXO), and with and without the positive allosteric modulator (PAM) LY2119620, were reported in Ref.^27^, where it was shown that LY2119620 is positioned to block the iperoxo egress route through the ECV. Later, several experimental studies of the binding kinetics of iperoxo with and without LY2119620 were performed ^28, 29^. Computationally, iperoxo dissociation was simulated using metadynamics by Capelli et.al.^30^ Two egress routes of iperoxo through the ECV were observed in these simulations with one being predominant. It was suggested that the presence of the PAM would block dissociation by both routes. In a subsequent study^31^, the same authors derived the dissociation rate of iperoxo using well-tempered-and frequency adaptive metadynamics to calculate the free energy profile for unbinding. It was concluded that further force field refinement might be needed to achieve an accurate transition-state free energy and thus unbinding rate constant ^31^. Interestingly, the binding kinetics of the radioligand, [^3^H]iperoxo, were observed to be bi-phasic in experiments in Ref.^28^, with two distinct characteristic times determined for both the dissociation and the association processes. Support for this biphasic behavior was not found in the theoretical study described in Ref. ^31^.

Here, we apply the τRAMD protocol to compute residence times for the alprenolol-β2AR complex and for three complexes of mAChR M2, with acetylcholine bound and with iperoxo bound in the absence and in the presence of LY2119620. We first analyze the egress routes observed in the RAMD trajectories in the individual proteins and compare them with previous simulation studies. For the analysis of the dissociation pathways and detection of metastable states, we employ MD-IFP for protein-ligand interaction fingerprint (IFP) analysis of dissociation trajectories^17^. We specifically investigate the influence of the single adjustable parameter in the τRAMD protocol, the random force magnitude, on the observed pathway distribution, in particular on the ligand egress between the transmembrane helices towards the membrane previously reported in some studies^32,33^, and on the unbinding mechanism, including the protein and ligand solvation processes during dissociation. Finally, we compare the relative τ values computed with the τRAMD protocol for all four complexes with available experimental data and find robust agreement between computation and experiment.

## 2. METHODS

The complete workflow for the τRAMD procedure and trajectory analysis is illustrated in Fig.1. The workflow includes the generation of several sets of MD simulations as well as the post-processing of trajectories as described in the following sections.

**Figure 1:**
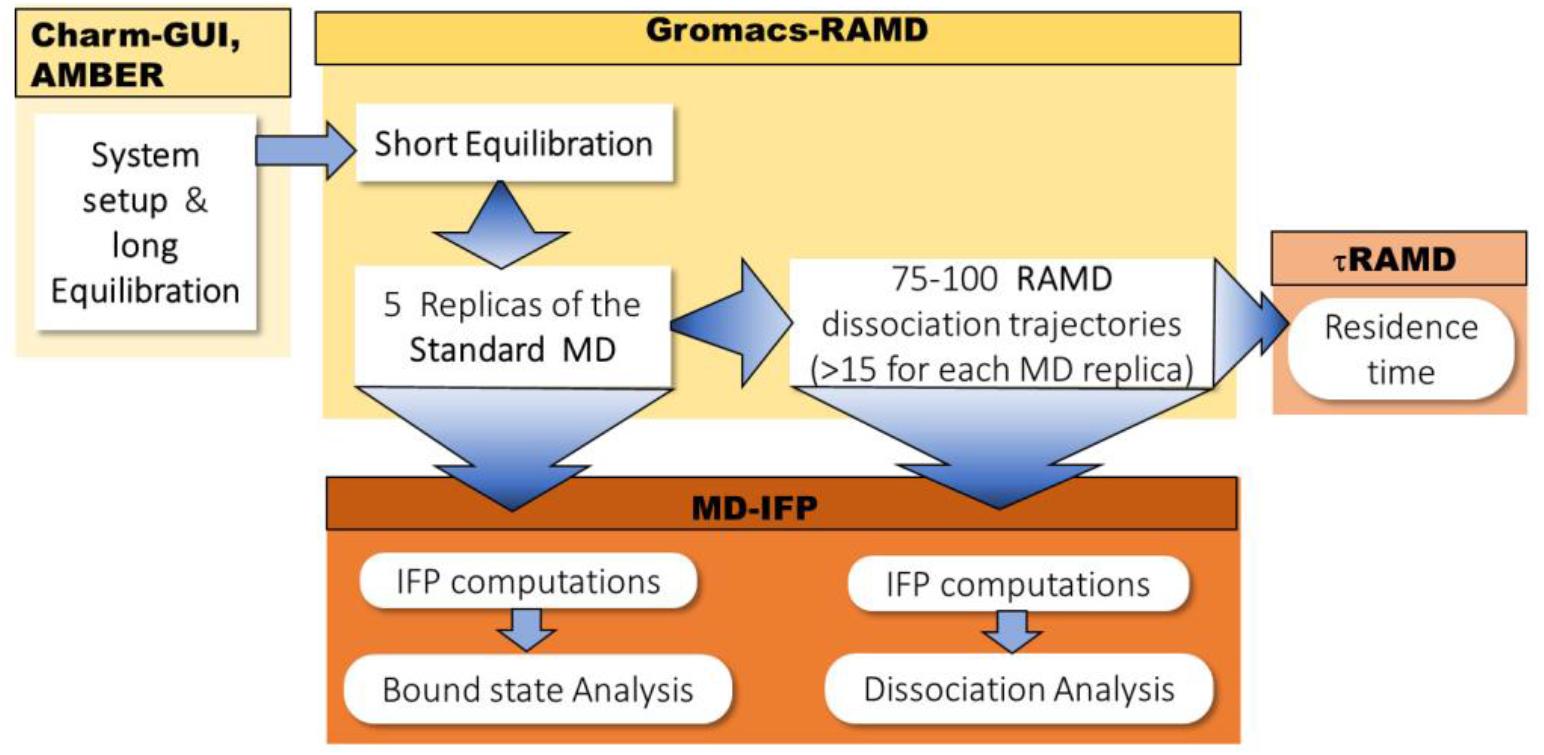
Schematic diagram of the workflow for τRAMD and MD-IFP analysis. The workflow includes a preparation step to set the systems up, conventional MD and RAMD simulations using the Gromacs-RAMD simulation engine, and trajectory analysis to compute relative residence times and a set of protein-ligand IFPs to explore the ligand dissociation profile (see main text). The shapes and sizes of the arrows approximately indicate the amounts of data generated in the individual steps.

### System setup and force field parameters

Complexes were built using the crystal structures with PDB IDs: 4MQT ^27^ (iperoxo agonist bound to the active conformation of mAChR M2 with allosteric compound LY2119620 solved at 3.70 Å resolution) and 3NYA^34^ (complex of the antagonist alprenolol with β2AR solved at 3.16 Å resolution). Protein residues are numbered according to the PDB files and additional superscripts give the numbering according to the GPCR secondary structure elements. Ligands were simulated in their protonated states (charge +1e). To simulate a complex of muscarinic receptor with iperoxo without LY2119620, we used the same crystal structure, 4MQT, for consistency, but removed LY2119620. The effect of the allosteric compound on the protein structure is minor, as shown by comparison with PDB ID 4MQS ^27^. For the simulation of ACh bound to mAChR M2, the iperoxo compound was replaced with ACh in the structure without LY2119620. In the simulations, we retained the engineered construct of GPCR with a fused intracellular domain that stabilizes the active conformation of the protein (a nanobody or T4 lysozyme for mAChR M2 and β2AR, respectively). Since RAMD simulations are relatively short (less than several tens of ns), large structural changes of the conformation of the protein (e.g. transitions from active to inactive conformations) are highly unlikely.

The CharmmGUI^35^ web server was used to setup systems with GPCR proteins embedded in a lipid bilayer by carrying out protein protonation at pH 7, and the generation of topology files and coordinates for simulations with the AMBER software package^36^. A complex membrane consisting of 50% cholesterol, 30% POPC (1-Palmitoyl-2-oleoyl-phosphatidylcholine), and 20% 1-palmitoyl-2-oleoylphosphatidylethanolamine (POPE) to represent the main components of a neuronal/synaptic membrane^37^ was built using the Amber Lipid14 force field^38^. The systems were solvated with TIP3P^39^ water molecules with a margin of at least 10 Å and Na^+^ and Cl^-^ ions were added to ensure system neutrality at an ion concentration of 150 mM. For simulations with ligands, the GAFF^40^ force field was employed. RESP partial atomic charges were computed for the ligands using GAMESS^41^ calculations of the electron density population at the HF/6-31G* (1D) level and Amber tools^42^.

### Equilibration MD simulation protocol

In all cases, for the first step of preparation, we employed the Amber software^36^, as in the previously published τRAMD protocol. One should note, however, that the entire protocol could be performed solely using the Gromacs software.

The system was first energy minimized (with restraints on all heavy atoms except water and ions of 1000, 500, 100, 50, 10, 1, 0.5, 0.1, 0.05, 0.01 kcal mol^-1^ Å^-2^ each for 1000 steps of conjugate gradient and then 10000 steps without restraints). It was then heated step-wise over 200ps (NVT-Langevin, tau = 1 ps^-1^) with restraints of 100 kcal mol^-1^ Å^-2^ up to 100 K. Then it was heated over 200 ps to 310 K with restraints of 5 kcal mol^-1^ Å^-2^. Following this, a short equilibration of 4 ns at 310 K with restraints of 5 kcal mol^-1^ Å^-2^ was performed. Then we removed the restraints and followed the protocol for the setup of simulations of membrane-containing systems on GPUs (https://ambermd.org/tutorials/advanced/tutorial16/) that consists of 10 consecutive simulations of 5 ns duration (separate simulations were used since the GPU code does not recalculate the non-bonded list cells during a simulation). Then we ran a further 300 ns of NPT (Langevin thermostat at 310K with a Berendsen barostat at 1bar) simulation to ensure equilibration of the whole system, including the membrane. For all simulations, a cutoff of 10 Å for nonbonded Coulombic and Lennard-Jones interactions and periodic boundary conditions with a Particle Mesh Ewald treatment of long-range Coulombic interactions were used. A 2 fs time step was employed with bonds to hydrogen atoms constrained using the SHAKE algorithm^43^.

Then the equilibrated systems were used in the GROMACS τRAMD protocols^17^ (for which a tutorial can be found at https://kbbox.h-its.org/). To perform simulations in GROMACS^44^, the final output coordinate and topology files were transferred from Amber to GROMACS using ParmEd^45^. Then we first performed short NVT simulations (Berendsen thermostat, 30 ns) and then generated 4-5 trajectories (**Table S1**) under NPT conditions (Nosé–Hoover thermostat and Parrinello-Rahman barostat, ∼20ns). Each trajectory was started with velocities generated from the Maxwell distribution to ensure trajectory diversity. The final coordinates and velocities were used for simulation of dissociation trajectories using RAMD (performed under the same NPT conditions).

### τRAMD protocol

The τRAMD protocol for computing relative residence times was reported in Refs. ^16,17^. Here, we briefly outline the main steps. A set of starting snapshots is generated from independent trajectories (from 4-6 replicas, **Table S1**) of conventional MD simulations as described above. The additional force is applied to the center of mass of an orthosteric ligand (except for a test run with the force applied to the PAM only). Each starting snapshot is then used to generate a series of at least 15 RAMD dissociation trajectories. We retained the default parameters of the RAMD protocol as described in detail elsewhere^17^ : the ligand displacement was evaluated every 100 fs, and a ligand displacement of less than 0.025 Å led to selection of a new random force orientation; unbinding was considered to have occurred when the ligand-protein COM separation distance reached 65 Å; the trajectory coordinates were recorded every 2 ps. The length of the RAMD trajectory was limited to 24h wall-clock time by the set-up of the compute cluster used. Within this time, about 55 ns could be simulated for the systems studied here. Correspondingly, the smallest possible external force magnitude was defined as 12 kcal/mol Å by the dissociation time of the slowest dissociating compound (around 45 ns for iperoxo from mAChR in the presence of PAM) for all ligands except ACh, for which the smallest force magnitude applied was 10 kcal/mol Å. We investigated the effect on residence times and dissociation mechanisms of varying the random force magnitude by running calculations with values of 12, 14 and 16 kcal/mol Å. All other parameters of the RAMD protocol were kept constant as their variation has no significant effect on the relative residence time, although they may change the absolute values of the computed residence times. ^16^

The effective residence time for each replica was defined by the computed dissociation time, corresponding to 50% of the cumulative distribution function (**Fig. S1**). A bootstrapping procedure (5000 rounds with 80% of samples selected randomly) was performed to obtain the residence time for each replica, τ_repl_, which should converge to a Gaussian-like distribution if the sampling is sufficient. The final relative residence time, τ_RAMD_, was defined as the mean τ_repl_ over all replicas.

Computed relative residence times were plotted for each random force used as a function of the experimental values on a logarithmic scale. The mean standard deviations of the computed residence times were computed as defined in the previously reported τRAMD protocol of Ref.^16^

### Dissociation trajectory analysis

Protein-ligand Interaction Fingerprints, PL-IFPs, and Protein-Ligand REsidue contacts, PL-REs, were generated for the last 700 frames of each RAMD dissociation trajectory with the snapshots being saved with a stride of 2 ps, discarding most of the frames in which the ligand retains the bound state position. The PL-IFPs included hydrogen bonds, aromatic interactions, water bridges, salt bridges and hydrophobic contacts, as specified in Ref.^17^. The PL-REs were defined as protein residues within a distance of 5.0 Å from any heavy atom of the ligand. Additionally, the number of water molecules in the first water shell around the ligand, the RMSD from the bound position of the ligand, and the ligand COM were computed for each snapshot. If a dissociation event was not observed because the simulation time allowed was exceeded, the trajectory was discarded from the analysis, see statistics in **Table S1**. The PL-IFPs and PL-REs for each snapshot were stored as binary vectors with 0/1 values for each individual contact (i.e., residue and type of interaction). The vectors generated from all trajectories for each compound and for each random force applied were then collected in a binary matrix.

To detect the most visited regions in the protein-ligand interaction space, we employed the PL-REs matrix as it is less sensitive to relative residue-ligand position and thus, provides more robust details of transient protein-ligand contacts of the complex, than more specific PL-IFPs. We employed kmeans clustering of the PL-REs matrix in the space of the contact residues using the corresponding function with default parameters as implemented in the scikit-learn package^46^. The selection of the number of clusters to be generated is the main uncertainty of the kmeans approach. In the present case, we had to make a trade-off between the difficulty of analyzing multiple clusters and the blurring of the protein-ligand contact specificity in the case of a small number of clusters. To assess consistency within clusters and obtain hints on the number of clusters to be selected, we employed the Silhouette method and selected 10 clusters, which is a compromise considering the high variation of Silhouette plots for the systems studied (**Fig. S2**). Additionally, we computed the contribution of the PL-IFPs to each cluster, which shows the population of specific protein-ligand interactions.

Different dissociation paths were distinguished using hierarchical clustering of the PL-REs vectors for the last protein-ligand contacts in each trajectory. Then, all trajectories assigned to one cluster have the same egress pathway (i.e., the same protein-ligand interactions in the last frame before complete unbinding), although ligand rotation and ligand motion inside the protein may vary.

For each snapshot, the positions of the ligand can be projected onto physical space by mapping the ligand COM onto a 3D grid. This enables the generation of a ligand COM density distribution of all of the selected RAMD dissociation trajectories and the superimposition of this density on the protein structure. Additionally, we generated ligand COM distributions for each cluster by summing over all snapshots assigned to the cluster. Importantly, snapshots assigned to a single cluster may not necessarily have a small RMSD relative to each other. Thus, the COM distribution in a cluster may not be compact or, alternatively, different ligand orientations with close COMs may be assigned to different clusters. The dissociated state is defined by a cluster in which no protein-ligand contact is found or in which only a few non-specific contacts are present, with the ligand COM spread around the protein. In contrast, the clusters describing the bound states of the ligands are usually compact in the physical space.

## 3. RESULTS AND DISCUSSION

### 3.1 Dissociation mechanisms of the alprenolol-β2AR complex

#### The main ligand egress routes are through the ECV whereas dissociation between the transmembrane helices is diminished upon reduction of the random force magnitude

Alprenolol binds in the deeply buried orthosteric cavity of β*2AR* (**Fig. 2)**, where it is coordinated by a salt bridge and a H-bond between its ammonium and hydroxyl groups and the N312^7.39^ and D113^3.32^ protein side-chains, respectively, and by an edge-to-plane aromatic interaction of its benzene ring with F290^6.52^. A direct exit tunnel through the extracellular vestibule, ECV, is partially blocked by the Y308^7.35^ and F193^ECL2^ side-chains dividing the cavity into two main regions in the crystal structure (see upper inset in **Fig. 2B**): one on the side of H5 and H6, and the other on the side of H2, H3 and H7. Additionally, a small sub-pocket is observed on the other side of the ECL2 loop (these three cavities are denoted by the three black arrows in Fig.2A). These cavities can be associated with different dissociation routes, as will be discussed below. The exit route from the ECV is covered in part by the ECL2 loop whose position is stabilized by the K305^7.32^-D192^ECL2^salt-bridge between ECL2 and H7.

**Figure 2:**
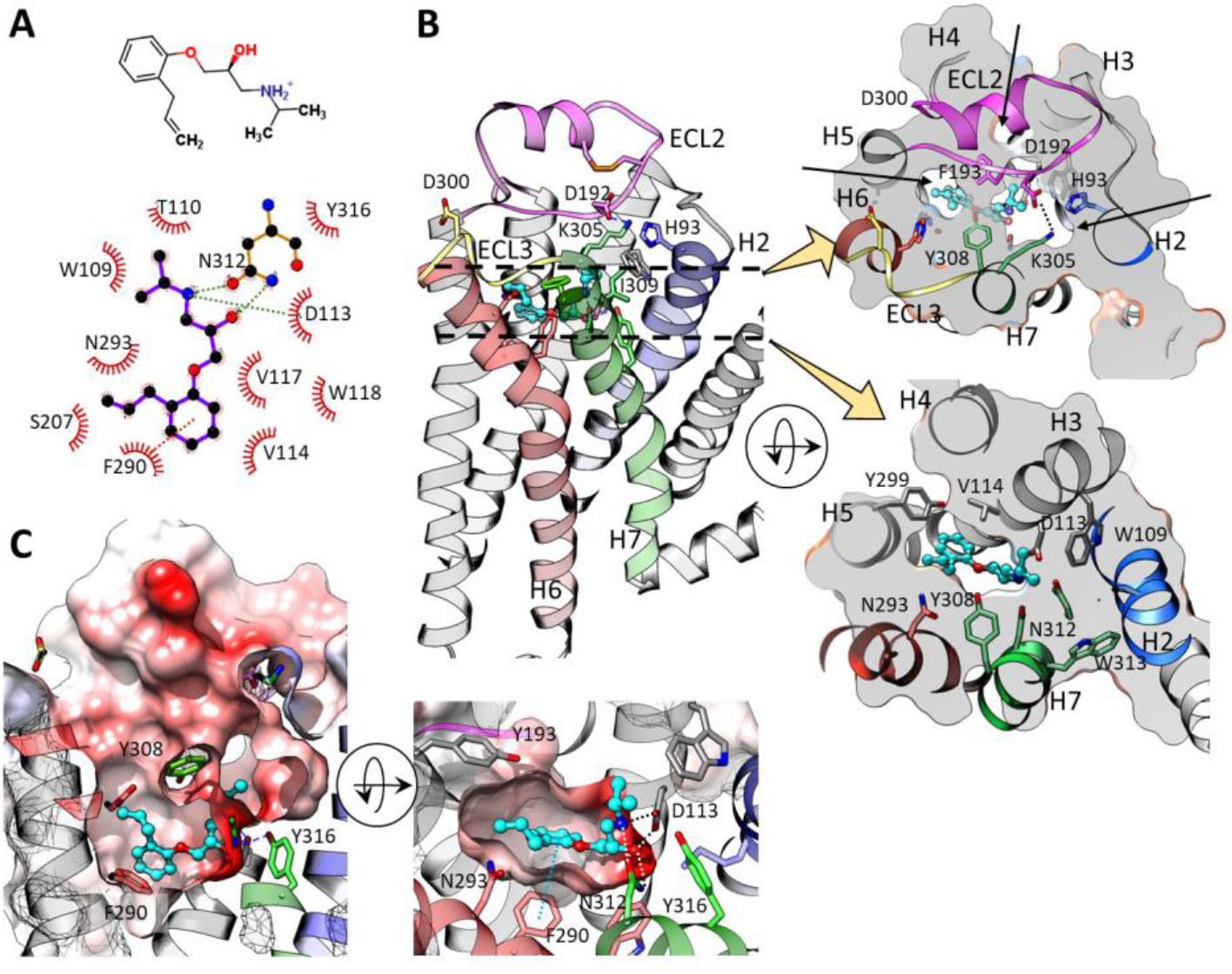
The structure of β2AR with alprenolol bound in the orthosteric site: (**A**) 2D representation of the ligand and protein-ligand interactions (generated using the Marvin Sketch ^47^and LigPlot+ ^48^ software, respectively); H-bonds and aromatic interactions are shown by green and brown dotted lines, respectively; (**B**) The bound structure (PDB ID 3NYA ^34^) is shown with the protein in cartoon representation with key loops and helices colored and selected binding site residues shown in stick representation. Alprenolol is shown in cyan carbon ball and stick representation along with its 2D structure. The shape of the cavity is shown in the insets by two cross-sections at the planes indicated by the black dashed lines. Upper inset: Cross-section through the protein surface cutting across the extracellular vestibule (ECV) roughly in the plane of residues D192^ECL2^, D300^ECL3^ and K305 ^7.32^, showing three subcavities (indicated by black arrows) extending from the ligand binding site to the protein exterior with the salt bridge D192 ^ECL2^-K305^7.32^ (dashed line) stabilizing the ECL2 loop and thereby closing the exit route from the ECV. Lower inset: Cross-section through the orthosteric binding pocket. **(C)** The orthosteric binding pocket is shown in two perpendicular views with the surface colored by Coulomb charge (red-white-blue pallet corresponds to negative to positive charge) and ligand-protein H-bonds in the complex shown by dashed lines.

In the RAMD simulations, alprenolol mainly leaves the pocket through the ECV via one of three partially overlapping dissociation routes (denoted ECV1, ECV2, and ECV3; see also next section) (**Fig. 3A**). Some trajectories are observed to lead into the membrane via several exit routes (**Table 1, Fig. 3A**). In the most populated routes into the membrane, the ligand passes through the narrow openings between helices H4 and H5 (path H4/5, blue arrow in **Fig. 3A**) that is also observed in the crystal structure (see **Fig. 2B**). The other pathways into the membrane are not readily apparent in the crystal structure and require protein conformational changes. The total occurrence of the egress routes into the membrane is dependent on the magnitude of the random force: about 32 % and 11% of the RAMD trajectories pass into the membrane in simulations with a magnitude of 16 kcal mol^-1^ Å^-1^ and 12 kcal mol^-1^ Å^-1^, respectively (**Table 1**). This indicates that the dissociation paths of alprenolol between the transmembrane helices in β*2AR* can be expected to become less likely in an unperturbed system. Indeed, in conventional MD simulations of the binding of alprenolol to β*2AR* ^24^, ligand partitioning from the aqueous environment into the lipid bilayer was observed, but alprenolol was not observed to enter the protein binding pocket from the lipid bilayer.

**Figure 3:**
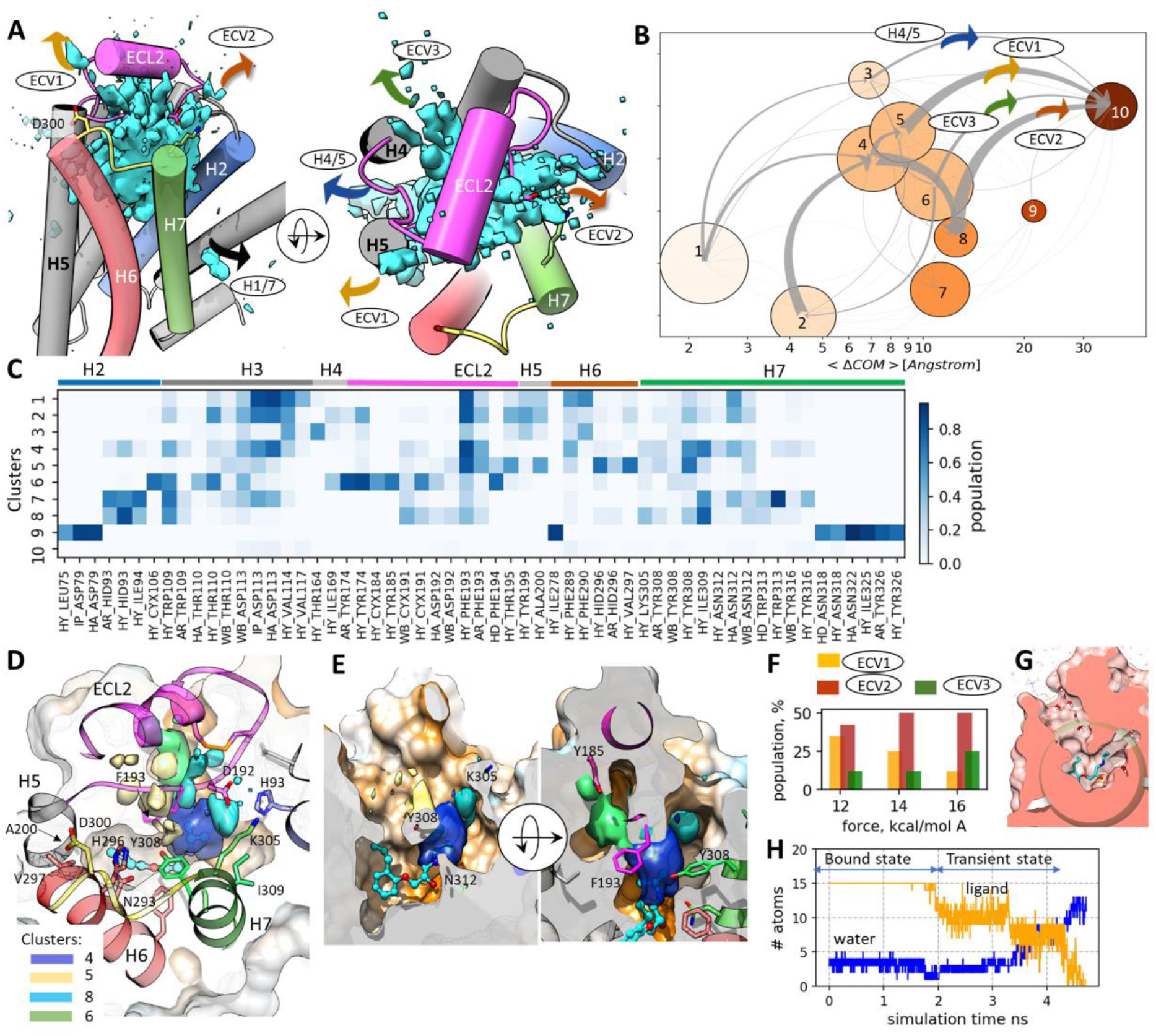
Ligand dissociation paths and their characteristics from RAMD simulations of the β2AR-alprenolol complex: (**A**) Side and top views of the relative ligand COM distribution in the RAMD dissociation trajectories with ligand pathways labelled (cyan iso-surfaces enclose regions visited more than once). (**B**) Ligand dissociation pathways displayed in a graph representation from PL-IFP analysis: each of the ten clusters is shown by a node with the size indicating the cluster population and positioned on an increasing logarithmic scale of the average ligand COM displacement in the cluster from the starting snapshot; the darker the node color the higher the average ligand RMSD in the cluster from the bound starting structure. The gray arrows indicate the total flow between two nodes. (**C**) IFP composition of the ten clusters which are ordered from 1 to 10 by increasing RMSD; the cluster composition for all ligand-protein contacts is shown in **Fig. S3**. (**D**) Top view of β2AR with superimposition of the metastable states (clusters 4, 5, 6 and 8) that are intermediates on the main dissociation pathways ECV1 and ECV2; metastable states are depicted as iso-surfaces of the ligand COM densities in the corresponding clusters. (**E**) Perpendicular views of the orthosteric pocket with the main metastable states colored as shown in (**D**); (**F**) Histogram showing the variation of the relative populations of the three main egress routes for different RAMD force magnitudes. (**G, H**) Solvation of the binding pocket along a RAMD trajectory: the number of water molecules was counted inside the sphere of radius 7 Å shown in (**G**) with a cross-section through the protein surface centered at the orthosteric binding pocket. (**H**) The number of water molecules (blue) that enter inside the pocket increases as the number of ligand carbon atoms (yellow) inside the pocket decreases. Results are shown for a representative RAMD trajectory simulated with a random force magnitude of 12 kcal mol^-1^ Å^-1^ (similar plots for further RAMD trajectories are displayed in Fig.S4).

**Table 1.** Egress statistics in the RAMD simulations and computed τRAMD τ values for the complexes with β2AR and mAChR M2

| GPCR | Ligand | % of trajectories showing ligand egress through the membrane<br>(a) | | | Computed $\tau$ , ns | | | Measured $\tau$ , s | Measured $K_D$, nM |
| --- | --- | --- | --- | --- | --- | --- | --- | --- | --- |
|  |  | Random force magnitude, kcal mol<sup>-1</sup> Å<sup>-1</sup> |  |  |  |  |  |  |  |
|  |  | 12 (10) | 14 | 16 | 12 (10) | 14 | 16 |  |  |
| β2AR | alprenolol | 11 | 25 | 32 | 13.9±1.2 | 4.32±0.9 | 1.14±0.5 | <21.7<sup>49</sup><br><260<sup>50(b)</sup> | 0.6 -2.<sup>51(c)</sup><br>7<sup>49</sup> |
| AChR M2 | ACh | 23 (18) | 18 | 28 | 1.88±0.66 (17.2±8.4) | 0.64±0.27 | 0.30±0.12 | ~43<sup>52(d)</sup><br>~1.1 s <sup>53(e)</sup> | 35.2±3.4<sup>52</sup><br>34.3±8.6<sup>29</sup> |
|  | iperoxo | 2 | 14 | 15 | 20.8±6.3 | 6.40±1.45 | 2.65±0.90 | ~97±18<sup>28(f)</sup><br>173±11 <sup>29(g)</sup> | 10.23±0.06<sup>28</sup><br>0.736 ±0.192<sup>29(h)</sup> |
|  | iperoxo with LY2119620 | 17 | 22 | 20 | > 55<sup>(i)</sup> | 15.3 ±5.6 | 3.62±2.54 | (3.75±1.20)<br>10<sup>3</sup> <sup>29(j)</sup> | 0.071±0.011 <sup>29(h)</sup> |

(a) Trajectories with at least one frame without protein-ligand contacts but with more than 3 ligand-membrane contacts were defined as egress routes through the membrane.
(b) $t_{1/2}$ < 15s and < 3min at 37°C in Ref. <sup>49</sup> and Ref. <sup>50</sup>, respectively.
(c) The smaller value was derived from $k_{on}$ and $k_{off}$ values and the larger value is from an equilibrium binding assay.
(d) Agonist competition assay ([<sup>3</sup>H]ACh); $k_{-1}$ ~1.38 min<sup>-1</sup> <sup>52</sup>.
(e) Washout of the agonists; using reversal of the FRET signals induced by atropine and methoctramine: 677 ± 68 ms and 609 ± 87 ms.
(f) [<sup>3</sup>H]iperoxo radioligand assay; the dissociation was described as biphasic with $t_{1/2}^{(1)} = 1.12 \pm 0.21$ min and $t_{1/2}^{(2)} = 54.6 \pm 6.46$ min, with the slow fraction constituting about 33%. We used an effective residence time estimated as $\tau = (1/\ln 2)(1/t_{1/2}^{(1)} + 0.33/t_{1/2}^{(2)})^{-1}$ .
(g) $k_{off} = 0.347 \pm 0.023$ min<sup>-1</sup>.
(h) EC<sub>50</sub> values; LY2119620 reduces iperoxo potency by about 11-fold.
(i) The maximum length of RAMD trajectories was limited to about 55 ns (average MD simulation length per day (24 hours)). Since complete ligand dissociation was observed in only 46% of trajectories, one can estimate that the computed τRAMD must be at least 55ns.

The present results can be compared with previous RAMD simulations of the β2AR-carazolol complex^23^. Carazolol has a similar size to alprenolol and an almost identical binding pose with similar contacts to the protein. In addition to egress routes to the ECV, dissociation between the transmembrane helices was recorded in about one third of the trajectories, which is comparable with the present simulations at the random force magnitude of 16 kcal mol^-1^ Å^-1^ (∼32% of trajectories), although the force applied in Ref.^23^ was almost an order of magnitude higher (acceleration 0.2-0.3 kcal g^-1^Å^-1^, corresponding to 60-90 kcal mol^-1^ Å^-1^). One should note however that, along with other differences in the parameterization of the calculations, a pure POPC bilayer was employed in Ref.^23^, in contrast to the complex membrane composition simulated in the present study, which may affect the flexibility of the transmembrane helices and membrane-ligand interactions.

#### RAMD simulations reveal the main features of the dissociation mechanisms observed in conventional MD simulations along with a broader distribution of dissociation pathways

The density distribution of the ligand COM in RAMD dissociation trajectories shown in **Fig. 3A** indicates that there is no single well-defined dissociation path. Instead, the distribution of ligand egress trajectories is quite diffuse. To explore the mechanism of ligand dissociation, namely the key protein-ligand interactions and metastable states, we analyzed the dissociation trajectories in the space of protein-ligand interactions. For this purpose, the last 700 frames of the RAMD trajectories were clustered by their ligand-protein residue contacts (PL-REs). Then, a transition matrix and the net flux between clusters were computed (see Methods section). The resultant graph of metastable states along the main egress routes of alprenolol from β*2AR* computed with the random force magnitude of 12 kcal mol^-1^ Å^-1^ is shown in **Fig. 3B** along with the PL-IFPs contributing to each cluster (**Fig. 3C**; the cluster composition in the PL-REs space and for the larger random force magnitudes is shown in **Fig. S3**).

Clusters 1 and 2 shown in **Fig. 3B** have RMSD < 5Å and comprise the starting ligand poses (note, that since only the last 700 frames from each trajectory were analyzed, the starting pose may deviate from the original bound state). They have an almost identical composition in the IFP space (see heatmap in **Fig. 3C**) with the main contacts being similar to those in the crystal structure: ammonium - salt bridge with D113^3.32^ and hydrogen bond with N312^7.39^ as well as hydrogen bonds of the ligand hydroxyl group with D113^3.32^, N312, and a set of hydrophobic contacts, such as the alprenolol phenyl ring with V114^3.33^, F193^ECL2^ and F290^6.52^. All other clusters, except for unbound-state cluster 10, represent intermediate metastable states.

According to the graph in **Fig. 3B**, the main ligand dissociation flow passes through the hub-like cluster 4, located around or immediately after the tunnel bottleneck lined by Y308^7.35^/ N312^7.39^, and leading to the metastable states 5 or 8 in the ECV (shown in **Fig. 3D, E**), and then proceeds to dissociation through the ECV (exit routes ECV1 and ECV2). While cluster 4 still bears some similarity to the bound state clusters, the transition to metastable states 5 and 8 occurs by the breaking of the hydrogen bonds between the ligand and D113^3.32^ and N312^7.39^, and the formation of new pathway-dependent hydrophobic and aromatic interactions, such as with Y308^7.35^/H296^6.58^/F193^ECL2^ and I309^7.36^/H93^2.47^, lining ECV1 and ECV2 pathways, respectively (**Fig. 3C**). Metastable states 8 and 5 are located under the D192^ECL2^-K305^7.32^ bridge and around F193^ECL2^, respectively (**Fig. 3D,E**), and represent the last metastable states prior to complete ligand unbinding through the ECV. The same is also true for metastable state 6, located in the cavity behind ECL2 (specifically, the F193^ECL2^ side chain) and leading to the egress route ECV3.

The trajectories leading to the ECV can be approximately separated into three partially overlapping egress routes: about 41% follow path ECV2 between H2, H7 and the end-part of ECL2, about 34% follow path ECV1 between H5 and H6, and about 12% emerge on the other side of ECL2, close to H4 and H3 (via clusters 8, 5 and 6, respectively). Clusters 3 and 9 lead to dissociation into the membrane between H4 and H5 or H2 and H7 (egress routes H4/5 and H2/7), for 8% and less than 1% of trajectories, respectively. Notably, as the random force magnitude is decreased, the relative population of routes ECV2 and ECV3 decreases, while the relative population of route ECV1 increases (see **Fig. 3F**).

The ligand dissociation mechanism observed in RAMD simulations can be compared with the ligand association mechanism reported in the conventional MD study of Ref.^24^, where two major association steps were identified: (i) ligand entrance into the ECV associated with ligand contacts with residues F193^ECL2^, H296^6.58^, V297^6.59^, A200^5.39^ and Y308^7.35^, and (ii) passage into the binding pocket, typically after a few hundred nanoseconds. This sequence of events can be traced in reverse in the RAMD simulations.

The first association step in the ECV is in excellent agreement with cluster 5 (see **Fig. 3C, D**). Thus, the path ECV1 through cluster 5 (yellow arrow in **Fig. 3A**) resembles the dominant route of alprenolol entrance into the binding pocket revealed in 11 out of the 12 successful ligand binding events recorded in conventional MD simulations ^24^. Interaction with D192 ^ECL2^, mentioned in Ref. ^24^, is more important to egress route ECV2, where alprenolol passes between ECL2 and helices H2 and H7, which was observed only once in the conventional MD simulations.

In the second association step reported in Ref. ^24^, alprenolol moves from the extracellular vestibule to the binding pose, forming a transient ammonium – D113^3.32^ salt bridge before a crystallographic contact with N312^7.39^ is formed. Similarly, the hydrogen bond to N312^7.39^ appears less often in the IFP analysis than the salt-link to D113^3.32^ during egress, (even in cluster 1 which deviates from the starting bound position as only the last 700 snapshots were included in the analysis).

Several spontaneous unbinding events were also recorded in the conventional MD simulations of Ref.^24^ They were described as having the ligand first moving from the binding pose into the ECV, then passing between Y308^7.35^ and F193^5.32^ towards solvent. Correspondingly, in the binding free energy profile computed later in metadynamics simulations in Ref. ^26^, two energy minima were found: a deep one, corresponding to the bound state, and a shallow local energy minimum located between two barriers to ligand exit (the first for the ligand passing between Y308^7.35^ and F193^5.32^, and the second for ligand solvation upon leaving the ECV). ^26^ This observation agrees qualitatively with the metastable states found in the present RAMD simulations. Indeed, we see the ligand transition from the bound state to a compact metastable state 4 located in the vicinity of these two residues (see **Fig. 3B, D**), and the next step is associated with the transition through the Y308^7.35^- F193^5.32^ gate towards more diffuse metastable states 5 and 8 located in the ECV followed by dissociation.

In the conventional MD study of Ref.^24^, it was concluded that the ligand solvation in the ECV is associated with an energy barrier. It is, therefore, interesting to see if similar effects are observed in RAMD simulations. To analyze the solvation process in RAMD trajectories, the water molecules and alprenolol carbon atoms inside the binding pocket (defined by a sphere of radius 7 Å around V114^3.33^, see **Fig. 3G**) were counted in the last 2000 snapshots. There are 15 carbon atoms of alprenolol and several water molecules inside the sphere when the ligand is in its bound position. During the ligand dissociation, the number of ligand atoms in the sphere decreases to zero at the end of the trajectories, while the number of water molecules increases, generally up to about 15 molecules. The time from the point when the ligand starts to move from its binding pose up to its complete departure from the pocket varies considerably from one trajectory to another: from several hundred picoseconds up to nanoseconds, as illustrated for a selected trajectory in **Fig. 3H** (see also **Fig. S4**). However, in general, the pocket solvation happens gradually from the moment the ligand starts to move out of the pocket. In conventional MD simulations of ligand binding, the replacement of the water molecules in the ECV by the ligand was observed to occur within a 1 ns ^24^, which is in good agreement with RAMD simulations when the random force magnitude is assigned a value small enough to ensure dissociation trajectory lengths on the nanosecond time scale. Interestingly, the ligand egress from the pocket often occurs step-wise (as illustrated in **Fig. 3H**), which may be associated with a solvation barrier, although RAMD simulations do not provide direct evidence for this.

Furthermore, in accord with conventional MD simulations^24^, we also observed breaking of the K305^7.32^-D192^ECL2^salt bridge and the opening of the gap between Y308^7.35^ and F193^5.32^ on the time-scale of a hundred picoseconds, which does not have any notable correlation with ligand unbinding and indicates that this motion is rather spontaneous and therefore cannot be rate-limiting for ligand unbinding kinetics. We did not observe any large-scale protein conformational changes. Although this may be due to the short simulation timescale, it also indicates that such large protein motions are not required for ligand dissociation; this is also consistent with the observations from conventional MD studies.

Despite the general similarity in the ligand unbinding profile in RAMD and unbiased MD simulations, there is an important difference in the first unbinding step. Specifically, in RAMD trajectories, the ligand mainly retains its orientation in the binding site, keeping the ammonium group pointed towards H2 and H7, whereas in conventional MD ^24^, the ligand rotates by almost 180° pivoting on the phenylethyl moiety when associating from the ECV region to the binding pose, which may enable the ligand to squeeze through the narrow tunnel between ECL2 and helices H5 and H7 (**Fig. 2C**). This rotation process was also associated with the ligand transition over the highest energy barrier on the dissociation pathway in a metadynamics study ^26^. While this difference may arise due to differences in the force fields used, another possible explanation is that this barrier has a mainly entropic character as the ligand needs to undergo conformational changes and adopt a specific orientation to pass through a narrow tunnel leading to the egress route ECV1. In RAMD simulations, due to the acceleration of the egress process, the ligand does not have time to find an appropriate orientation. Instead, due to the additional random force, it is able to perturb Y308^7.35^ and F193^5.32^ and leave along a route requiring less orientational adjustment. The latter egress mechanism results in a higher fraction of egress events through pathway ECV2. Consistently, as the RAMD force magnitude is decreased and the dissociation time becomes longer, the flow through ECV2 is diminished in favor of the ECV1 route.

### 3.2 Dissociation mechanisms of agonists from mAChR M2

#### Egress routes are similar for both iperoxo and acetylcholine but change in the presence of the positive allosteric modulator

The binding mode of the orthosteric agonist iperoxo in mAChR M2 was reported in Ref.^27^ Two protein-ligand interactions, an H-bond between N404^6.52^ and the isoxazoline moiety and an ionic interaction of D103^3.32^ with the trimethylammonium moiety of iperoxo, coordinate the ligand in the plane almost perpendicular to the M2 transmembrane channel (**Fig. 4**). The aromatic rings of Y403^6.51^ and Y426^7.39^ also contribute to cation-π interactions with the positively charged trimethylammonium moiety of the ligand whereas the isoxazoline moiety makes hydrophobic contacts with V111 ^3.40^. ACh has a similar but less stable position (not shown). The protein channel is blocked towards the extracellular region by a lid formed by three tyrosine side-chains (Y104^3.33^, Y403^6.51^ and Y430^7.43^, **Fig. 4B**) interconnected by hydrogen bonds. Dissociation of iperoxo to the extracellular region therefore requires breaking the hydrogen bond network of the lid and flipping the tyrosine side-chains aside, or squeezing the ligand through the narrow channel that is lined by W155^4.57^ and connects the orthosteric binding site with the ECV (**Fig. 4C**). The positive allosteric modulator (PAM), LY2119620, bound at the top of the ECV (PDB ID 4MQT, see **Fig. 4A,B**), further blocks the egress of the orthosteric ligand, increasing the residence time of iperoxo by over tenfold in experiments^29^ (**Table 1**). Notably, LY2119620 does not affect the structure of the orthosteric binding pocket or the binding pose of the orthosteric ligand, and only slightly shifts the extracellular ECL2 loop and causes re-orientation of W422^7.35^, which forms a pi-stacking interaction with LY2119620^27^.

**Figure 4.**
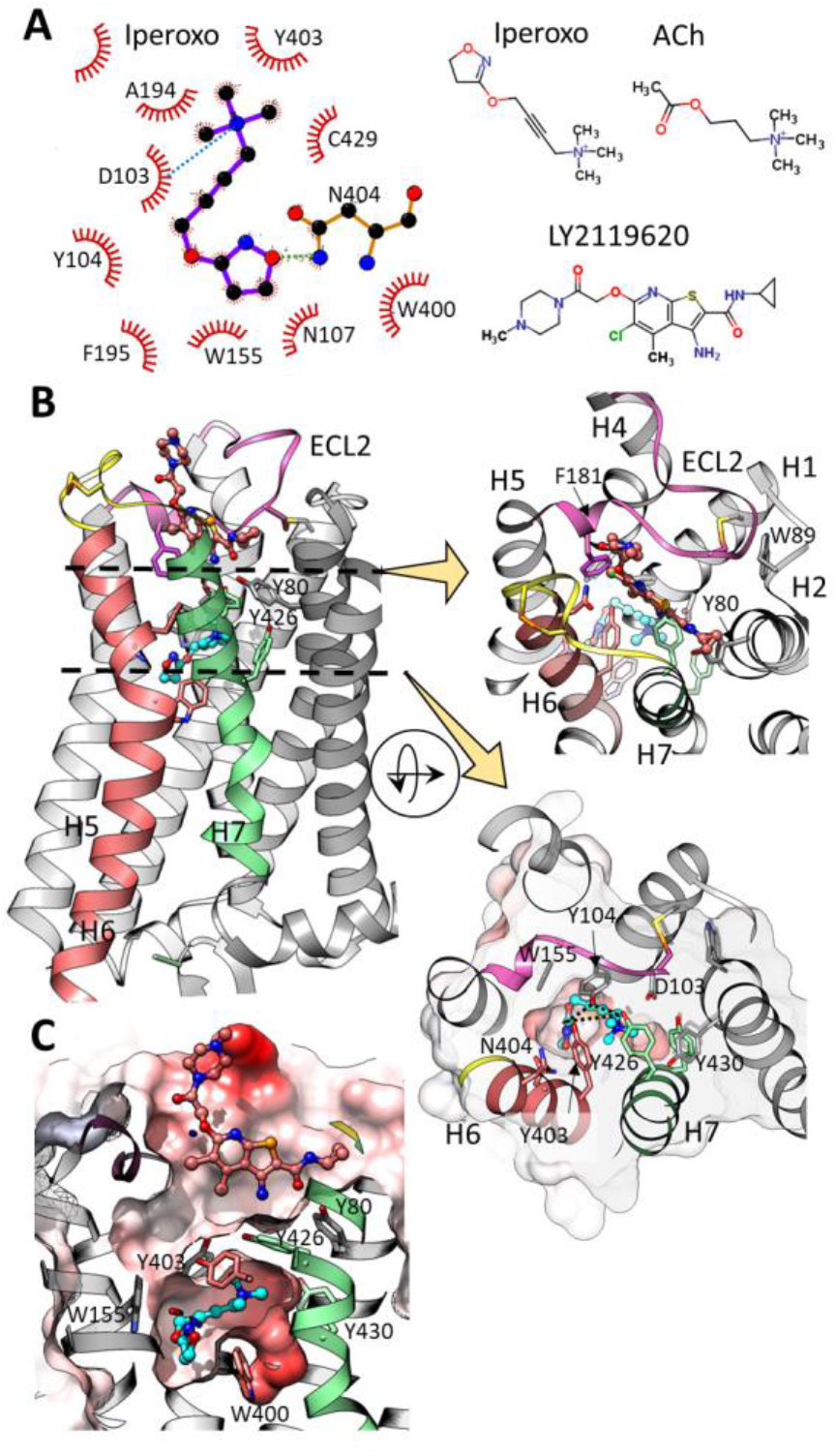
Structures of the complexes of mAChR M2 with iperoxo, the positive allosteric modulator (PAM), LY2119620, and acetylcholine (ACh), PDB ID: 4MQT. **(A)** 2D structures of the ligands and protein-ligand interactions for iperoxo (generated using the Marvin Sketch^47^ and LigPlot+^48^ software, respectively), H-bonds and ionic interactions are shown by green and blue dotted lines, respectively. **(B)** Structure of the M2 muscarinic receptor in complex with iperoxo and LY2119620 (PDB ID 4MQT, solved at 3.7 Å resolution^27^). The carbon atoms of iperoxo and PAM are colored cyan and salmon, respectively. The ECL2 extracellular loop is shown in pink and helices H6 and H7 are colored red and green, respectively. In the bound state, the iperoxo egress route is blocked by the side-chains of three tyrosine residues (Y104^3.33^, Y403^6.51^ and Y426^7.39^) linked to each other by hydrogen bonds, and by the allosteric compound positioned in the ECV. The two inset close-up views show the ECV with the PAM bound (upper) and the orthosteric pocket (lower). Protein-iperoxo H-bonds and H-bonds between the three tyrosines of the tyrosine lid are shown by black dashed lines in the lower inset. **(C)** The shape of the orthosteric and allosteric pockets is shown by the protein surface colored by Coulomb charge (red: negative; blue: positive); the narrow egress route from the orthosteric pocket is lined by W155^4.57^ and the tyrosine lid.

As for β*2AR*, the main egress routes in the RAMD simulations were through the ECV (ECV1-3) and the ligand passes between the transmembrane helices and into the membrane in a smaller number of trajectories (**Fig. 5A-C**). Among them, the two most populated paths run between H5 and H6 and between H4 and H5 (H5/6 and H4/5). As the magnitude of the random force is reduced from 16 to 12 kcal mol^-1^ Å^-1^, and the dissociation time reaches ∼10-30 ns, the fraction of iperoxo dissociation events into the membrane decreases from 15% to 2% (see **Table 1** and **Fig. S5**), similarly to β2AR. For ACh, however, this tendency is less pronounced: even at the random force magnitude of 10 kcal mol^-1^ Å^-1^ and a dissociation time above 10 ns, about 18% of egress pathways are still towards the membrane. Similarly, in the presence of the PAM, about 17% of ligand egress events are through the membrane, independently of the force applied, although the iperoxo dissociation time becomes several fold longer (**Table 1**). Apart from this, the PAM alters the dissociation pathways through the ECV: instead of ECV1, a new exit path, ECV3, through the region between H4, H5 and ECL2 emerges; the ECV2 route is still observed, albeit with some shifting towards H3 (**Fig. 5B**).

**Figure 5:**
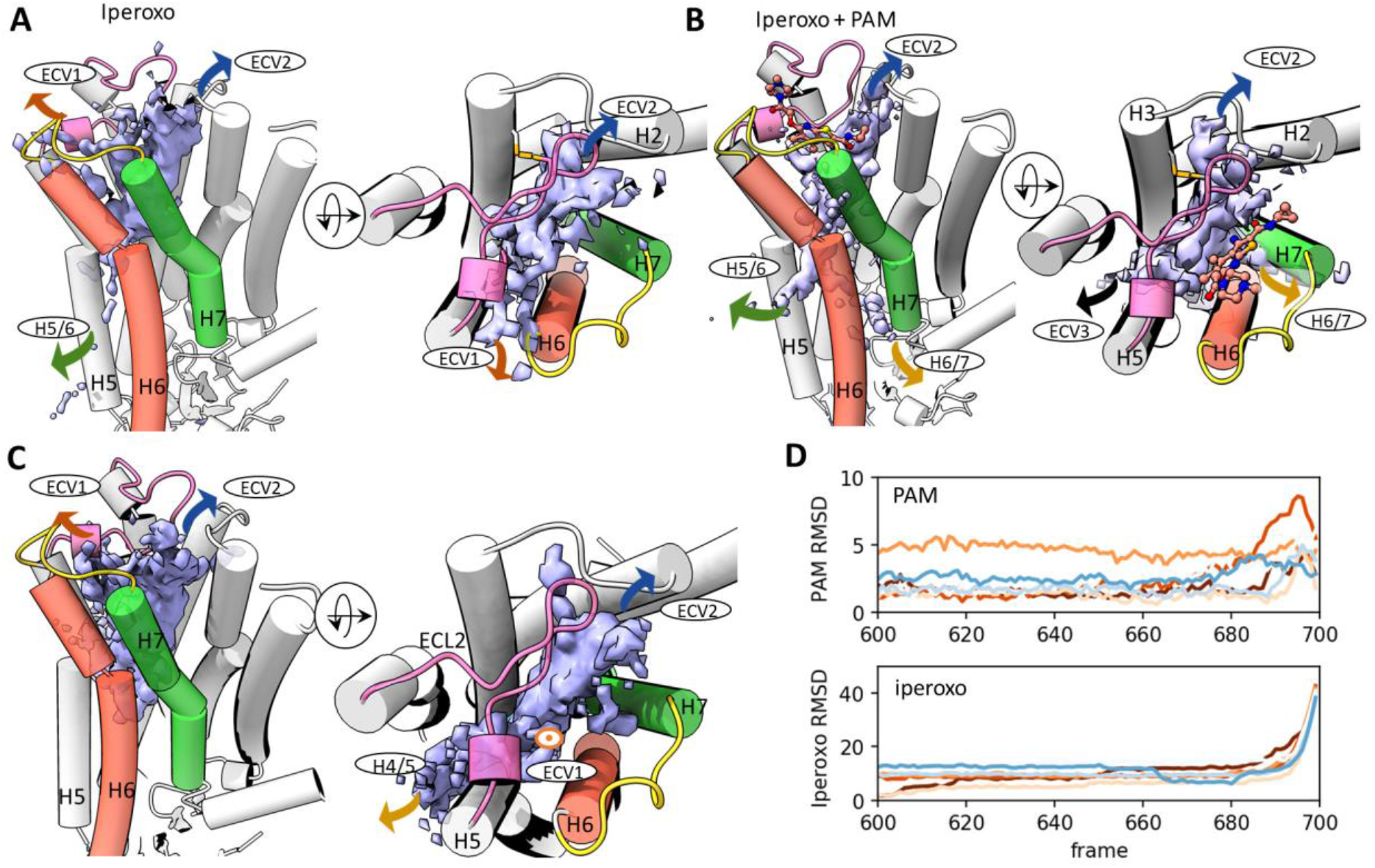
RAMD simulations of the egress of the iperoxo and acetylcholine agonists from the orthosteric binding site of mAChR M2. **(A-C)** Dissociation routes (indicated by arrows (or and ellipse if the direction is perpendicular to the figure plane) and labels) are shown by the ligand COM density isosurfaces contoured at 0.1 for the orthosteric ligand obtained in RAMD simulations with the minimum random force magnitude employed: 12 kcal mol^-1^ Å^-1^ for iperoxo **(A)**, and iperoxo with the PAM **(B)**, and 10 kcal mol^-1^ Å^-1^ for ACh **(C)** (see **Fig. S5** for dissociation routes at different forces). **(D)** Variation of the RMSD relative to the starting structure of the PAM and iperoxo (upper and lower plots, respectively) with time over the last 100 frames of RAMD trajectories. Each trajectory is shown in a different color and only trajectories in which the PAM RMSD reached a value greater than 4 Å are shown.

One might expect that prior to iperoxo dissociation, the PAM should leave the ECV. Surprisingly, this is not the case. In only about 20% of trajectories does the PAM completely or partially change its position and this occurs only in the final 100-200 picoseconds of dissociation (the RMSD of the PAM for several such trajectories is shown in **Fig. 5D**). On the other hand, the PAM dissociates very fast if the RAMD force is applied directly to it (τ_RAMD_ < 100 ps at a RAMD force magnitude of 12 kcal mol^-1^ Å^-1^).

#### RAMD simulations show similar dissociation mechanisms for the mAChR M2 agonists to those observed for the alprenolol-β2AR complex

Metastable states obtained from clustering of the last 700 frames of the dissociation trajectories generated with the smallest random force magnitude employed are shown for three mAChR M2 systems in **Fig. 6 A-C** along with the PL-IFP composition of each cluster (PL-RE cluster compositions for all systems are shown in **Fig. S6**).

**Figure 6:**
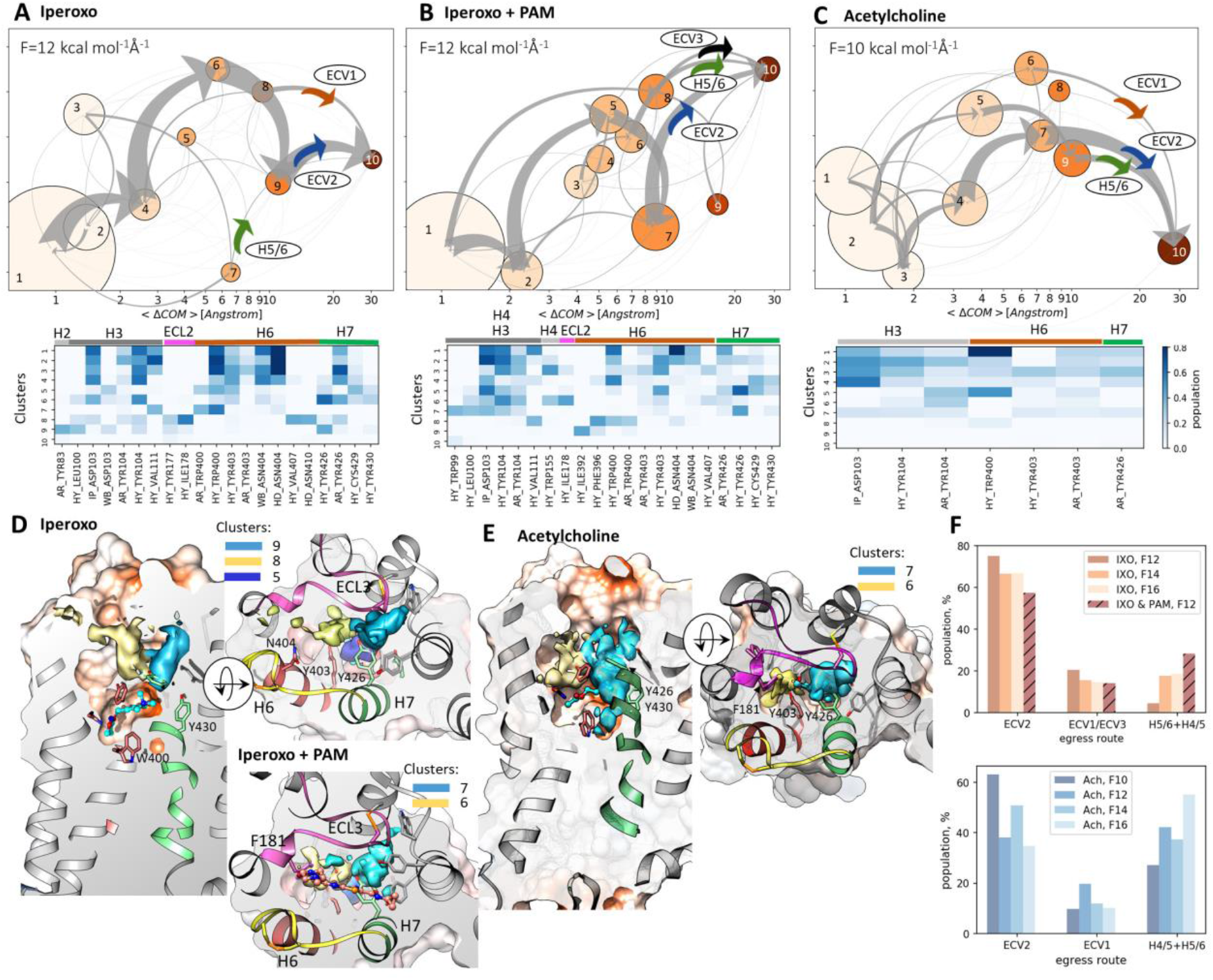
Analysis of RAMD dissociation trajectories of the ACh and iperoxo agonists from the orthosteric binding site of mAChR M2: **(A-C)** Dissociation pathways in a graph representation (same representation as for Fig. 2) with the corresponding IFP composition of each of the ten clusters shown as a heat map; **(D-E)** –metastable states leading to dissociation shown as COM distributions for each complex studied (Representation as described in legend to Fig. 3). **(F)** Fraction of trajectories along the most populated egress routes for iperoxo (upper histogram) and ACh (lower histogram) obtained from computing the flow to the unbound state for trajectories obtained with four different magnitudes of the random force.

Clusters corresponding to the ligand located in the orthosteric binding pocket (RMSD < 2Å ; clusters 1-3 / 1-2 for iperoxo without/with PAM; clusters 1-3 for ACh) are characterized by several main ligand binding contacts: a charge interaction between D103 and the positively charged trimethylammonium group of iperoxo/ACh, either cation-π or hydrophobic interactions of the tyrosine lid of Y104^3.33^ and Y403^6.51^ (and Y426^7.39^ for iperoxo only) with the trimethylammonium moiety, a hydrogen bond or water bridge between N404^6.52^ and the isoxazoline moiety of iperoxo, and hydrophobic interactions of V111^3.40^ and W400^6.48^ with the carbon atoms of the iperoxo isoxazoline ring.

There are multiple intermediate metastable states in which the orthosteric ligand is shifted from its starting position by over 3 Å (**Fig.6 A-C**). Among them, there are three relatively highly populated states that are directly connected with the unbound state (i.e., cluster 10) and represent the main intermediate steps before dissociation: two of them (8,9 / 6,7 for iperoxo without/with PAM, respectively, and 6,7 for ACh) are in the ECV and lead mainly towards the egress routes ECV1/ECV2 for iperoxo and ACh, and ECV2/ECV3 for iperoxo with PAM. The ECV2 pathway starts from the metastable state between ECL2, H2 and H7 (shown in light blue in **Fig. 6 D, E**), while pathway ECV1/ECV3 originates from the more diffuse metastable state separated from ECV2 by Y104^3.33^ and Y426^7.39^ and lined by F181^ECL2^ and Y403^6.51^ (shown in yellow in **Fig. 6 D, E**). All these metastable states are located around the tyrosine lid in the ECV. Thus, the general pattern of ligand dissociation resembles that for the alprenolol - β*2AR* complex, where a well-populated metastable state is located immediately before the aromatic side-chain gate and several less localized metastable states in the ECV.

#### Comparison of simulations reveals ligand egress features that are sensitive to the random force magnitude

Metadynamics simulations of the iperoxo-mAChR complex^30^ revealed that the first step on the ligand dissociation route was ligand rotation around the trimethylammonium moiety, which was locked by polar interactions with D103^3.32^ and the tyrosine lid, while breaking the hydrogen bond with N404^6.52^. This suggests some similarity between the two systems studied, iperoxo-mAChR M2 and β2AR-alprenolol.

In RAMD simulations of the iperoxo-mAChR complex, we observe similar, but not exactly the same ligand behavior. Indeed, the first step in the iperoxo dissociation profile is the loss of the interaction between the isoxazoline moiety and the protein (hydrogen bond with N404^6.52^ and hydrophobic contacts with V111^3.40^), while preserving the interactions of the trimethylammonium group with D103^3.32^ and Y104^3.33^ (cluster 4 for iperoxo and clusters 2 and 5 for iperoxo with the PAM). Note that although the tyrosine residues may interact with either the trimethylammonium group or the isoxazoline ring, the charge-charge interaction with D103^3.32^ is only possible with the positively charged trimethylammonium group, which suggests that at the first step, the protein-ligand hydrogen bond with the isoxazoline fragment breaks, while the charge interaction with the trimethylammonium group is preserved. Despite this, ligand rotation around the trimethylammonium-D103^3.32^ axis by more than 90° is observed only rarely in RAMD trajectories.

Moreover, the metadynamics simulations suggest that the ligand then follows the narrow channel lined by W155^4.57^, Y104^3.33^ and Y403^6.51^, which resembles the pathway ECV1 more than the most common route, ECV2, in the present simulations. According to Ref. ^30^, this motion is assisted by rearrangement of the ECL2 extracellular loop in metadynamics simulations, which is, not so strongly pronounced in RAMD simulations (the RMSD of the ECL2 heavy atoms remains within 3Å in all dissociation trajectories, likely due to the stabilization effect of the disulfide bridge to H1, which was absent in the simulations of Ref. ^30^; see **Fig. 4B**. It was suggested in Ref. ^30^, that a change in the position of ECL2 might enable opening of an additional egress route on the other side of ECL2. Although we do not observe sizeable rearrangements of the ECL2 position, the additional egress route ECV3, as well as displacement of the ECV2 channel, are indeed observed in RAMD simulations of iperoxo in the presence of PAM, as discussed above (see also **Fig. 5B**).

Comparing the dissociation mechanism of iperoxo from mAChR for different random force magnitudes with the corresponding simulations of the alprenolol-β*2AR* complex, we observe some similarities. In both cases, both conventional MD and metadynamics simulations reveal ligand rotation, which is needed to squeeze through a narrow channel, whereas such rotation of the ligand is rarely observed in the RAMD simulations. Instead, some side-chains that hinder direct exit are nudged away. It is important to note here that, in other systems where the ligand is less tightly constrained by the surrounding protein, interactions of the ligand with the protein due to the random force can facilitate significant ligand rotation ^19^.

Decreasing the force magnitude led to a change of the pathway population in favor of the routes requiring a stricter ligand orientation and revealed in conventional MD and metadynamics simulations: the relative populations of ECV1 to ECV2 changed on lowering the force magnitude from 16 to 12 kcal mol^-1^ Å^-1^ from 0.22 to 0.27 and from 0.24 to 0.83 for iperoxo and alprenolol, respectively (see **Fig. 3F** and, less pronounced, in **Fig. 6F**). However, no clear trend was observed for ACh.

### 3.3 Computed relative residence times correctly rank the four compounds for the two GPCRs

In **Fig. 7**, the computed residence times, τ_RAMD,_ are plotted versus measured residence times for all systems studied and for the three random force magnitudes applied. Remarkably, the relative values computed for all the different systems show a good linear correlation on a logarithmic scale with the experimental residence time for all forces applied (R^2^ = 0.94, 0.93, and 0.89 and the mean standard deviation for computed residence times is 30%, 29%, and 29% for the random force magnitudes of 12, 14 and 16 kcal mol-1 Å^-1^, respectively), which indicates the general applicability of the method. However, calculations with further complexes would be necessary to assess the performance across the GPCR superfamily.

**Figure 7:**
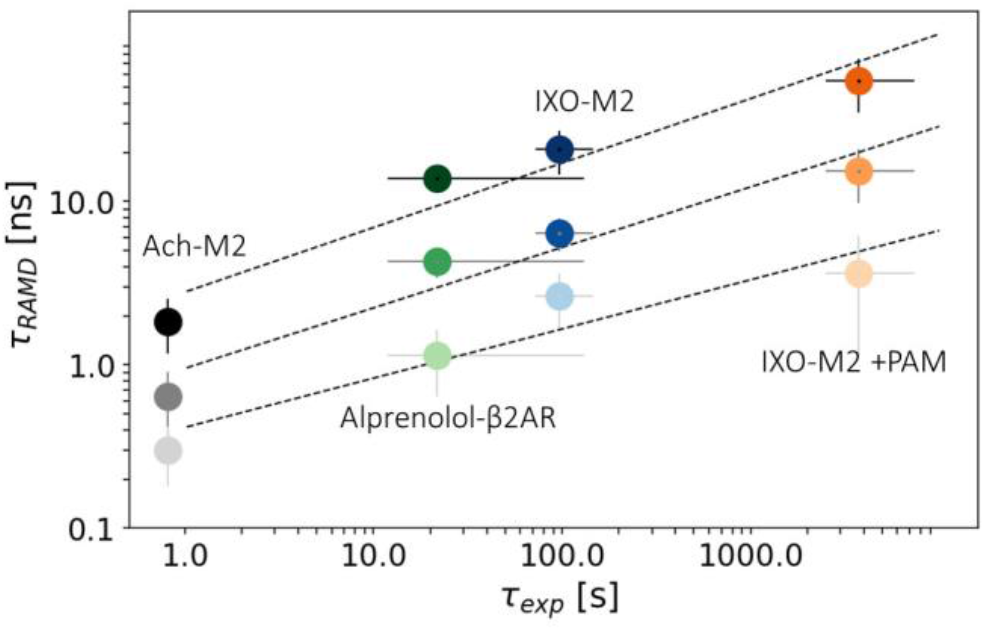
Computed relative residence times for the four complexes studied are correlated with the experimental residence times at all random force magnitudes. (see Table 1). The darkness of the color decreases with increasing random force magnitude from 12 kcal mol^-1^ Å^-1^ to 14 kcal mol^-1^ Å^-1^to 16 kcal mol^-1^ Å^-1^ with dashed lines showing linear fits of the computed values to the experimental ones. Note that for iperoxo (IXO) with PAM, the lower estimate of the computed residence time at 12 kcal mol^-1^ Å^-1^ is shown (see comments to Table 1). Experimental data are as given in Table 1.

The diversity of the ligand dissociation pathways is generally reduced for the smaller magnitudes of the random force. This could be expected to affect τ_RAMD_, for example, because egress routes to the membrane, if artifacts, may lead to some underestimation of the cumulative τ_RAMD_. As an example, we considered iperoxo in the presence of the PAM: the egress through the H5/6 route (**Fig. 5B**) is about 50% faster than through the ECV2 route (**Fig. S7**). From populations and residence times for each route shown in **Fig. S7**, one can estimate that if all transitions would occur through the ECV only, the residence time would increase by about 30%. This value may be larger in the case of ACh, where the fraction of ligand egress to the membrane is even higher than for iperoxo with the PAM.

Interestingly, iperoxo unbinding as observed in a radioligand [^3^H]iperoxo assay was best fitted by two exponential functions with t_1/2_ = 1.12 min and 54.6 min, with a slow fraction of about 33.5% Ref.^28^. In our simulations, the distribution of ligand dissociation times is quite broad but smooth (see **Fig. S8**) and lacks biphasic behavior. We also did not observe a large difference in dissociation times for different egress routes. Generally, it is unlikely to identify a biphasic unbinding rate in a study of complex dissociation using trajectories with limited length, since if several concurrent dissociation pathways with strongly different residence times exist, all dissociation events will tend to follow the fastest one.

## 3. CONCLUSIONS

We have compared the ligand unbinding mechanisms and kinetics for four GPCR complexes with orthosteric binders: alprenolol – β2AR and acetylcholine- and iperoxo-mAChR M2, as well as iperoxo in the presence of the positive allosteric modulator, LY2119620. For the simulation of ligand unbinding and computation of dissociation rates, we have employed the τRAMD protocol, based on random acceleration MD simulations, where an additional force is applied to the ligand center of mass to enhance ligand dissociation from minutes or hours to the nanosecond timescale.

The main egress routes and protein-ligand interactions in the key metastable states were found to closely resemble those obtained from long conventional or metadynamics MD simulations, though RAMD dissociation pathways are more spatially delocalized and, in some cases, their relative populations are different. Analysis of the dissociation trajectories showed that the unbinding pathways of both the slow-dissociating agonists, iperoxo and alprenolol, are characterized by two main metastable states: one defined by protein residues blocking the exit channels and the second one located in the ECV and likely associated with the ligand solvation process. Dissociation of the smaller ACh ligand occurs through similar intermediate states, but these are less localized and therefore less stable, consistent with the shorter dissociation times observed in simulations and experiment.

We also observed that the variety of the dissociation paths diminished upon decreasing the magnitude of the random force applied, as did the fraction of dissociation events between the transmembrane helices toward the lipid bilayer in most systems. Examination of the dependence on the random force magnitude of the ligand egress routes and mechanisms provided a way to assess the robustness of the results and the relative preferences for the different egress routes observed.

We found that the dependence of the egress route population on the random force magnitude varied for the different systems studied. For iperoxo, which has an experimental residence time of several minutes, a force of 12 kcal/molÅ was required for ligand dissociation on the timescale of about 20 nanoseconds, which occurred predominantly through the ECV with hardly any egress to the membrane. In the presence of the PAM, however, the same random force magnitude leads to dissociation times over 55ns, with 17% of the egress events being into the membrane. Similarly, for ACh, despite the quite fast dissociation (for example, ∼2 ns for a random force magnitude of 12 kcal/molÅ), we did not observe a notable reduction of the fraction of dissociation events into the membrane, which was around 20% for all the random force magnitudes investigated. This indicates that dissociation into the membrane cannot be ruled out. Indeed, egress through the membrane has been suggested to be the major route in some systems (see, for example Ref.^54^). Besides, the membrane composition may also affect ligand egress to the membrane.

Remarkably, despite the diversity in dissociation path distribution in the RAMD simulations, the ranking of residence times is in a very good agreement with experimental data. The accuracy of the ranking is only slightly improved as the force magnitude decreases, which suggests that minor egress routes do not significantly affect the estimated unbinding rate. Importantly, even at the smallest force magnitude used in the present simulations, the accumulated total MD simulation time was in the range of 0.4 -1.5µs per ligand. This is still a much shorter simulation time than necessary for conventional MD simulations, which typically require about an order of magnitude longer time to observe a single binding event. Unbinding simulations of the ligands studied (with residence times on the seconds to minutes timescale) are currently not feasible at all without additional bias. Dissociation simulations using metadynamics require simulations of the timescale of several microseconds, but also strongly depend on the correct choice of collective variables, while collective variables do not need to be defined for RAMD simulations.

Finally, this study showed that dissociation of both mAChR M2 agonists, iperoxo and ACh, occurs along similar egress routes through the ECV. Binding of the positive allosteric modulator completely blocks one of the possible dissociation routes of iperoxo, leaving another partially open and additionally opening a new one. Although the present simulations do not allow the extrapolation of the ligand dissociation profile to the unbiased limit, the increase of the iperoxo residence time due to the PAM is in a good agreement with experimental data. This suggests that the τRAMD approach is applicable for the evaluation of the effect of allosteric modulators on the unbinding kinetics of orthosteric compounds.

## SUPPLEMENTARY MATERIAL

### Supplementary Figures

***Figure S1*** *Processing of the results of RAMD simulations with the τRAMD protocol for the estimation of the relative residence times for four complexes*

***Figure S2*** *Variation of the Silhouette score for different numbers of clusters (k on the x-axis) for clustering of RAMD trajectories for two systems and two different random force magnitudes*

***Figure S3*** *Dissociation profiles of alprenolol from* β*2AR obtained from RAMD simulations performed with different random force magnitudes*

***Figure S4*** *Illustration of the solvation of the* β*2AR binding pocket upon ligand dissociation in several RAMD trajectories computed with two random force magnitudes*.

***Figure S5*** *Two perpendicular views of the distribution of the ligand COM for three mAChR M2 complexes from RAMD simulations performed with different magnitudes of the random force*

***Figure S6*** *PL-REs populations of the clusters for three complexes of mAChR M2 obtained from RAMD trajectories generated with the minimum random force magnitude employed*

***Figure S7*** *Dissociation routes of iperoxo in the presence of a PAM. (A) Distribution of the ligand COM in RAMD trajectories colored by egress path*

***Figure S8*** *Dissociation times of iperoxo in all simulated RAMD trajectories computed with a random force magnitude of 14 kcal mol*^*-1*^ *Å*^*-1*^

***Figure S9*** *Dissociation of iperoxo towards the membrane between helices 4 and 5 in a RAMD simulation in the presence of the PAM, showing how the lipid is displaced by the ligand as it exits from the protein*

## Supplementary Tables

***Table 1*** *Statistics for the RAMD simulations*.

## Supplementary Movie

*A representative dissociation trajectory for the βA2R-alprenolol system*

This information is available free of charge via the Internet at http://pubs.acs.org

## DATA AVAILABILITY STATEMENT

Scripts and files for running the simulations, several representative RAMD trajectories, computed interaction fingerprints and Jupyter Notebooks for their analysis can be found at Zenodo https://zenodo.org/record/5151217#.YRASdI4zYuU (DOI: 10.5281/zenodo.5151217).

## AUTHORS’ CONTRIBUTIONS

DKB: Conceptualization, Methodology, Software, Validation, Formal analysis, Investigation, Data Curation, Writing – original draft, Writing – review & editing, Visualization.

RCW: Conceptualization, Writing – review & editing, Supervision, Funding acquisition

## ACKNOWLEDGEMENTS

We thank the European Union’s Horizon 2020 Framework Programme for Research and Innovation under Grant Agreements 785907 and 945539 (Human Brain Project SGA2 and SGA3), and the Klaus Tschira Foundation for financial support. We thank Riccardo Capelli and Paolo Carloni for fruitful discussions, Bernd Doser for the RAMD implementation in Gromacs, Stefan Richter for technical support, and Ariane Nunes-Alves and Giulia D’Arrigo for helpful comments on the manuscript.

