## Supplementary Tables and Figures for "G Protein-Coupled Receptor-Ligand Dissociation Rates and Mechanisms from τRAMD Simulations"


### I. Supplementary Tables

***Table 1*** *Statistics for the RAMD simulations. The maximum simulation time of a RAMD trajectory was limited to 55 ns due to the limits of the computer queuing system. In some trajectories, dissociation was not observed within this time limit. These trajectories were not considered in the further analysis to compute residence times.*

| Protein | Ligand | % dissociated | | | | Total no. of Trajectories (No. of Replicas) | | | |
|---|---|---|---|---|---|---|---|---|---|
| | | Random force magnitude, kcal mol$^{-1}$ Å$^{-1}$ | | | | | | | |
| | | 10 | 12 | 14 | 16 | 10 | 12 | 14 | 16 |
| *β2AR* | alprenolol | - | 83 | 89 | 100 | - | 75(5) | 75(5) | 60(4) |
| mAChR M2 | ACh | 85 | 95 | 97 | 100 | 75(5) | 70(4) | 75(5) | 60(4) |
| | iperoxo | - | 81 | 87 | 100 | - | 75(5) | 75(5) | 75(5) |
| | iperoxo in the presence of LY2119620 | - | 46 | 82 | 100 | - | 75(5) | 90(6) | 75(5) |

### II. Supplementary Figures

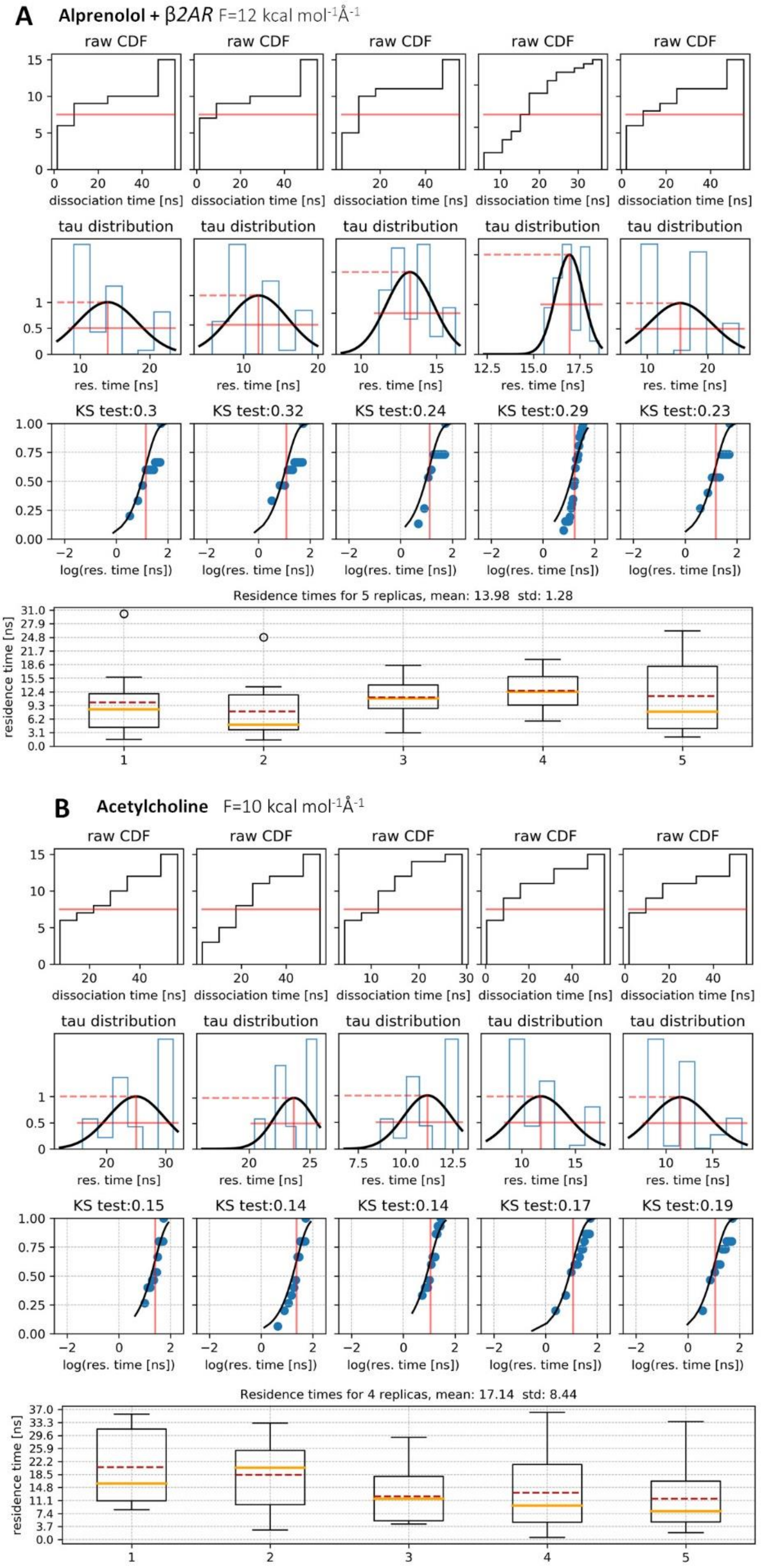

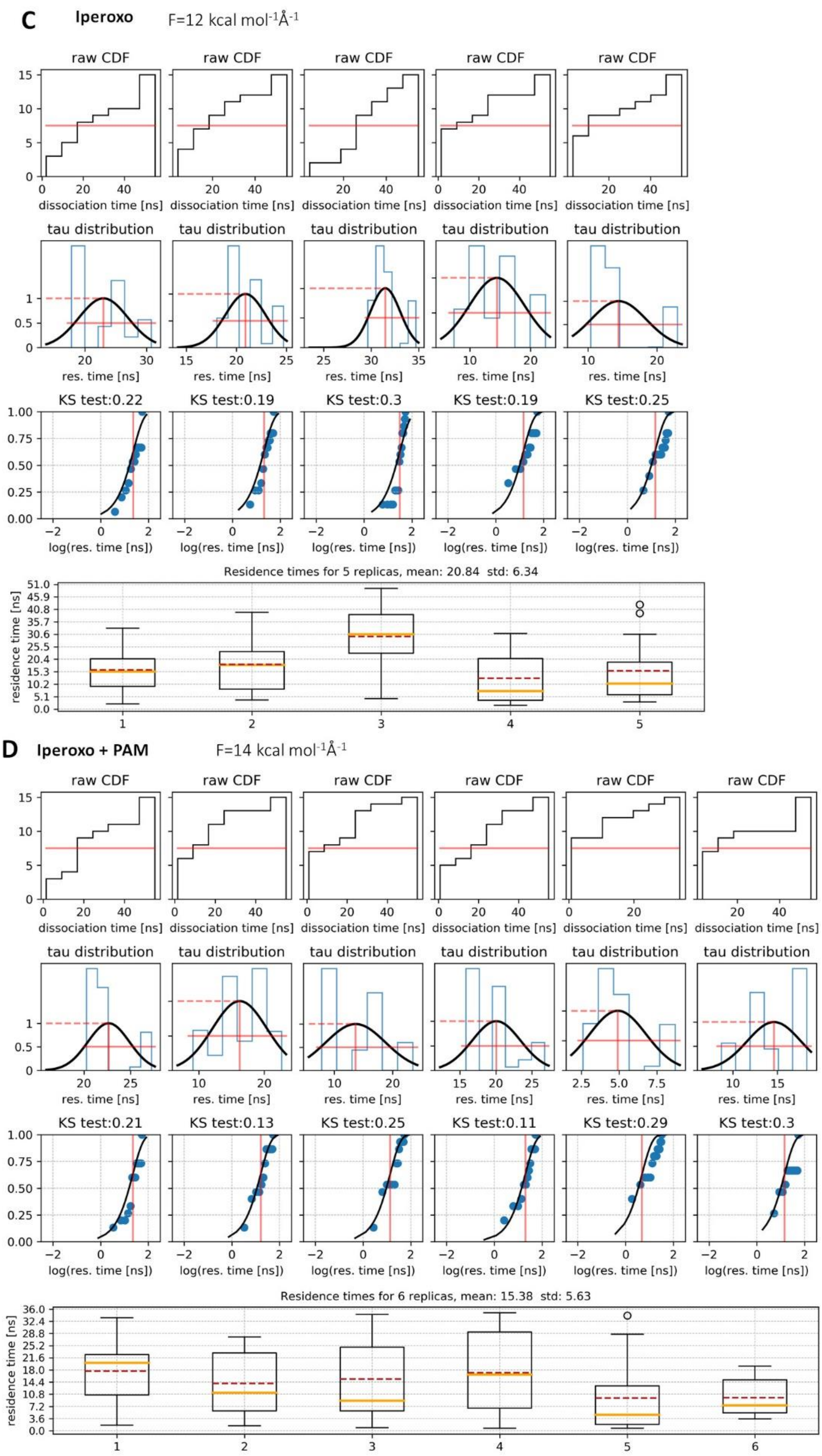


***Figure S1 Processing of the results of RAMD simulations with the τRAMD protocol for the estimation of the relative residence times for four complexes:*** *Alprenolol with β2AR and acetylcholine, iperoxo, and iperoxo in the presence of a PAM with mAChR M2. The following plots are shown for each complex: (1) Cumulative distribution function (CDF) for sets of RAMD trajectories*

*generated from five or six different replicas. The effective residence time obtained from these raw data is indicated by the red solid line at the time when the ligand has dissociated in half of the trajectories (1/2 of the maximum value of the CDF); (2) Distribution of effective residence times obtained after bootstrapping of the raw data along with the corresponding Gaussian distribution (black line); the mean and the half-width of each distribution are indicated by red lines. (3) Comparison of the Poisson cumulative distribution function (PCDF, black line) with the empirical cumulative density function (ECDF, blue points) obtained from the dissociation probability distribution. The residence time for each replica is indicated by the red line; the results of the Kolmogorov-Smirnov (KS) test are quantified by the supremum of the distance D between the Poisson and empirical CDFs (denoted above the plot); (4) Box plots of the relative residence times obtained for each replica. The box extends from the lower to the upper quartile values of the data, the whiskers show the range of the data, outliers are shown by black circles, the median and mean are shown by orange and dashed red lines, respectively. The average residence time in ns computed from all replicas is shown with its standard deviation above the box plot. From the figures, it can be seen that the residence times computed for each replica deviate from the mean residence time with the standard deviation value of up to ~15% for alprenolol and up to ~50% for ACh and for Iperoxo with PAM.*

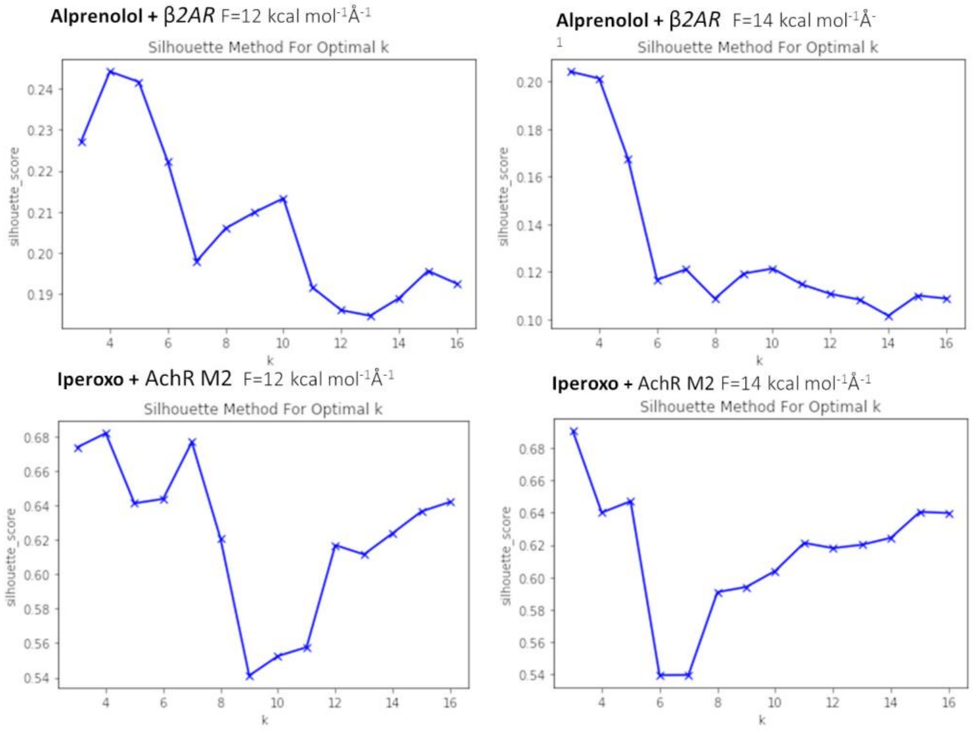


***Figure S2 Variation of the Silhouette score for different numbers of clusters (k on the x-axis) for clustering of RAMD trajectories for two systems and two different random force magnitudes.*** *Clustering was done using the PL-REs values extracted from the last 700 frames of each trajectory (see Methods section in the main text). The number of clusters (k) was chosen as 10 for the analysis of all trajectories as this value gave reasonably low silhouette scores in all cases.*

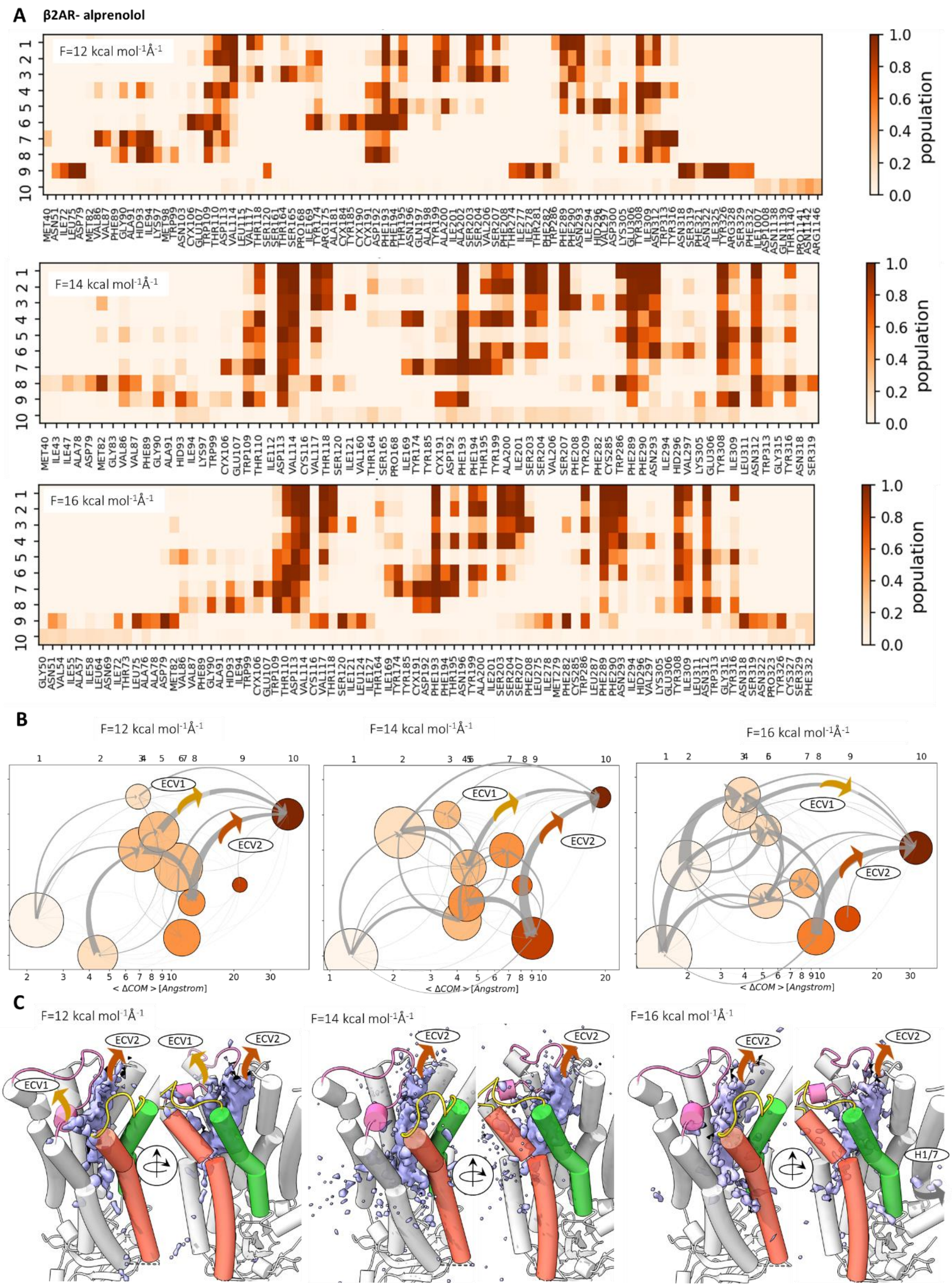


***Figure S3 Dissociation profiles of alprenolol from* β*2AR obtained from RAMD simulations performed with different random force magnitudes****: (A) IFP PL-REs composition of the ten clusters which are ordered from 1 to 10 by increasing RMSD (B) Graph representations of the metastable states and dissociation pathways; clusters are numbered by increasing <ΔCOM> distance and the*

*corresponding cluster numbering is given above each plot. (C) Side views of the ligand COM population density (lilac iso-surfaces) in the RAMD dissociation trajectories with ligand pathways labelled. See legend to Figure 2 in the main text for details.*

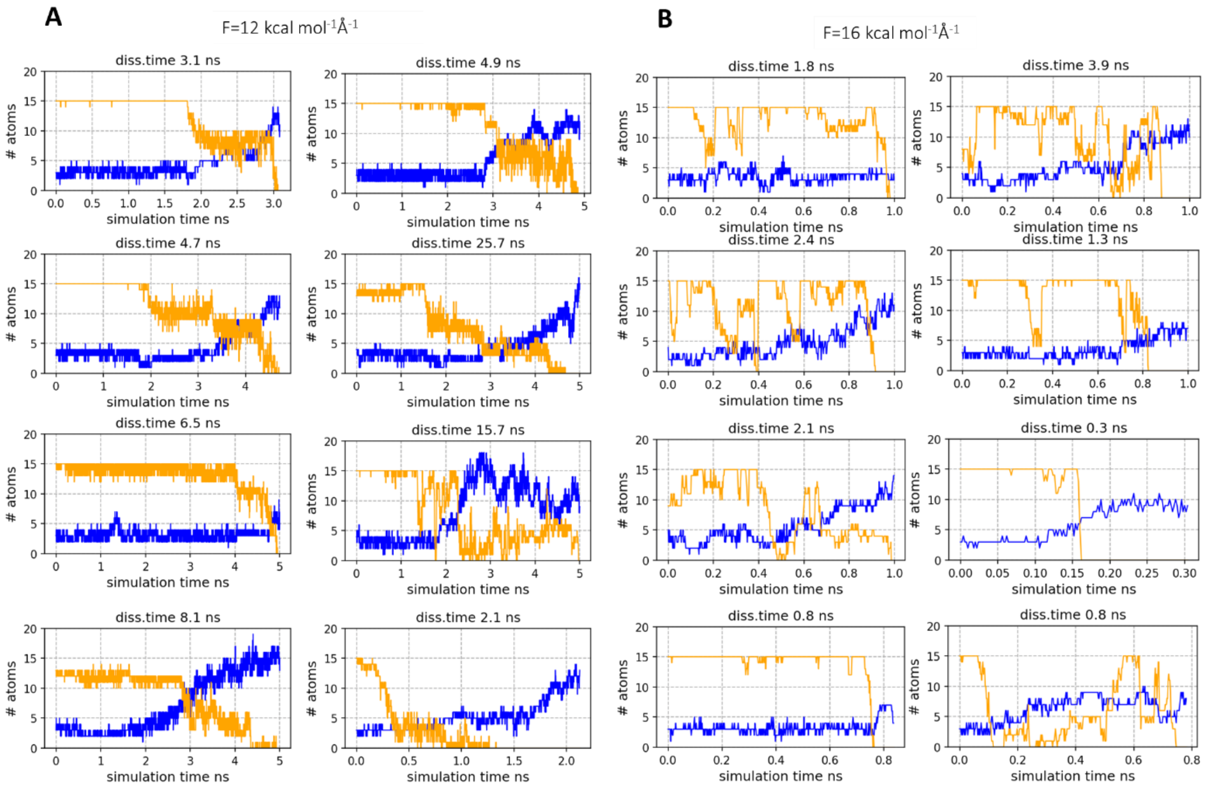


***Figure S4 Illustration of the solvation of the β2AR binding pocket upon ligand dissociation in several RAMD trajectories computed with two random force magnitudes.*** *The number of water molecules (blue) that enter inside the pocket increases and the number of ligand carbon atoms (orange) inside the pocket decreases as a function of time (see caption of Fig. 2G in the main text for more details).*

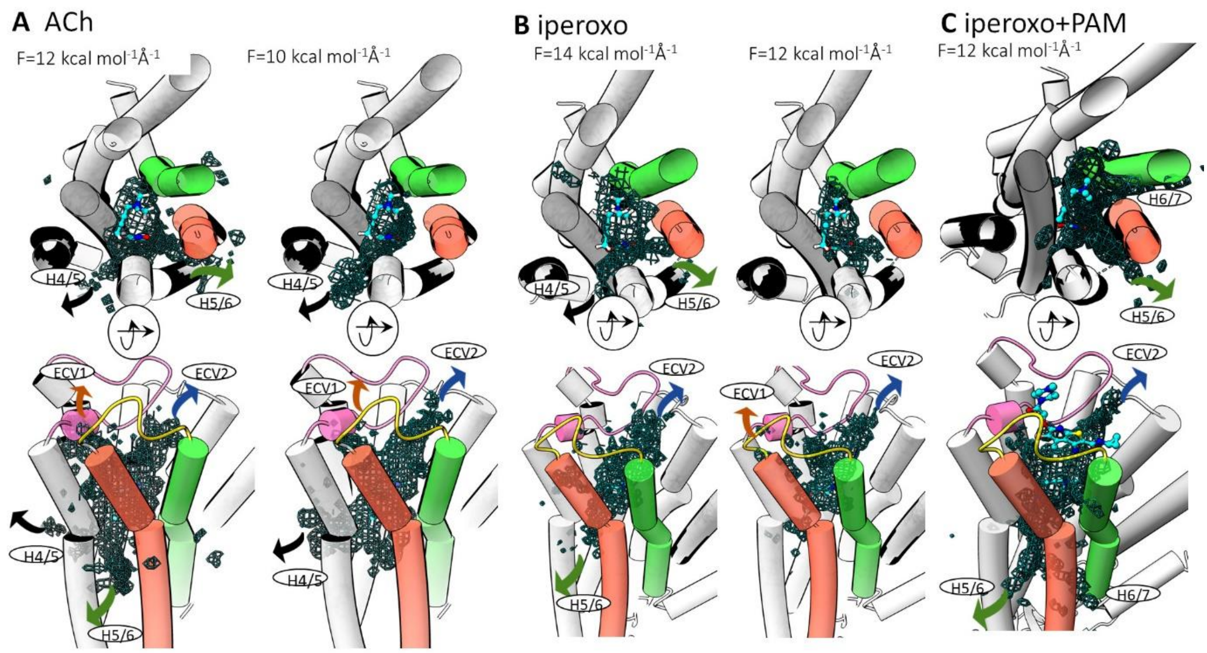


***Figure S5 Two perpendicular views of the distribution of the ligand COM for three mAChR M2 complexes*** *from RAMD simulations performed with different magnitudes of the random force. Egress routes are labelled. See legend of Figure 4 of the main text for details.*

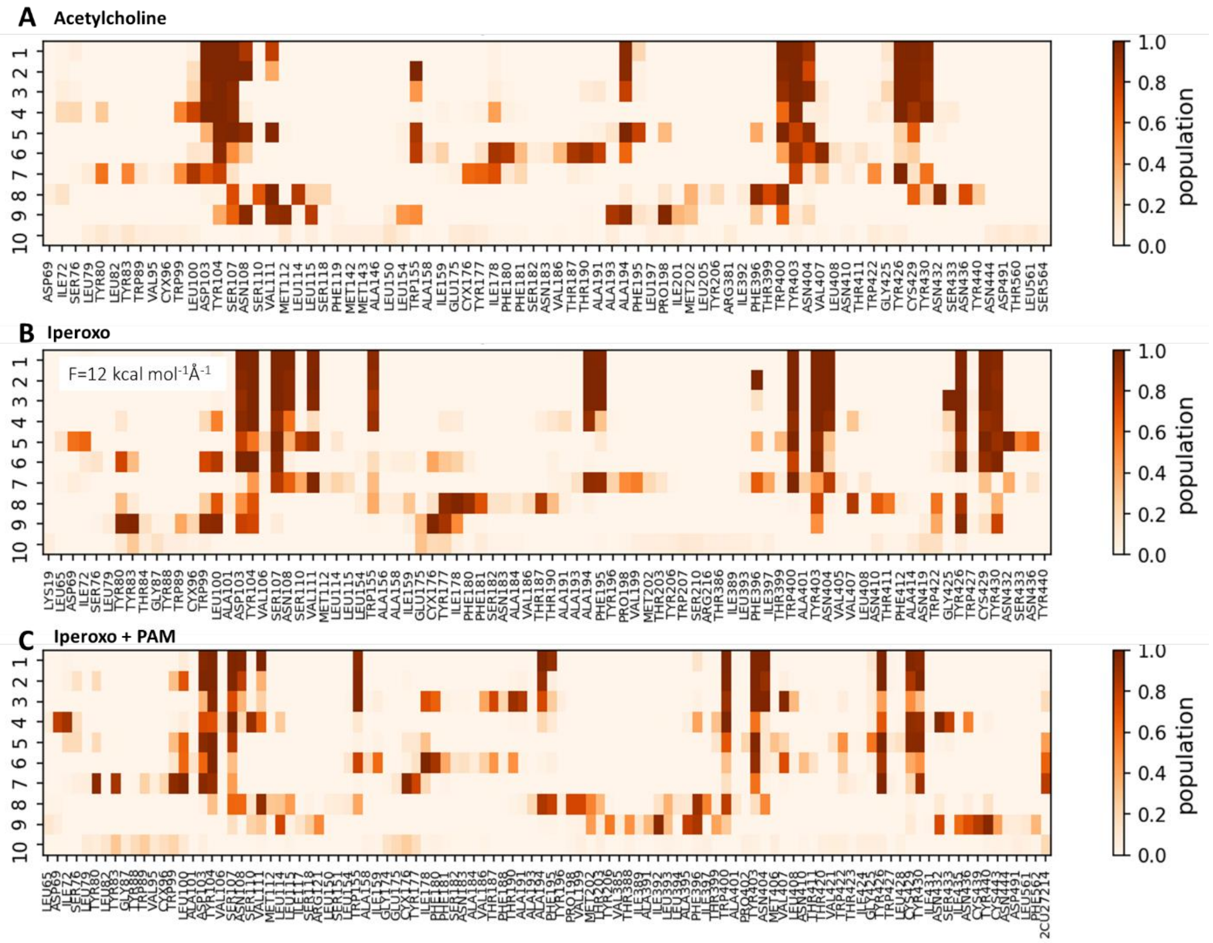


***Figure S6 PL-REs populations of the clusters for three complexes of mAChR M2*** *obtained from RAMD trajectories generated with the minimum random force magnitude employed: 10 kcal mol$^{-1}$ Å$^{-1}$ for ACh **(A)**, 12 kcal mol$^{-1}$ Å$^{-1}$ for iperoxo and iperoxo with PAM **(B)**, and **(C)**, respectively.*

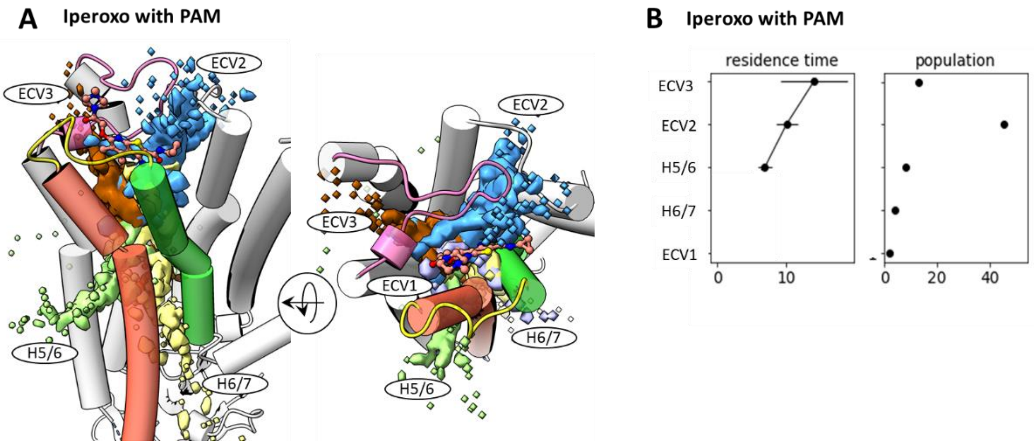


***Figure S7*** *Dissociation routes of iperoxo in the presence of a PAM. (A) Distribution of the ligand COM in RAMD trajectories colored by egress path (hierarchical clustering was used to split the trajectories into paths; see Methods section of the main text). (B) Residence times (ns) and populations (number of trajectories; the total number of trajectories resulting in dissociation was 74, see Table S1) for the five egress routes shown in (A), residence times was not computed for the two egress routes with very low populations (H6/7 and ECV1); the RAMD force magnitude was 14 kcal mol$^{-1}$ Å$^{-1}$.*

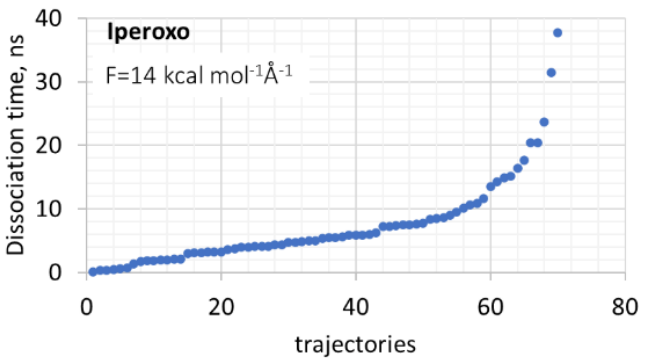


***Figure S8 Dissociation times of iperoxo in all simulated RAMD trajectories** computed with a random force magnitude of 14 kcal mol$^{-1}$ Å$^{-1}$.*

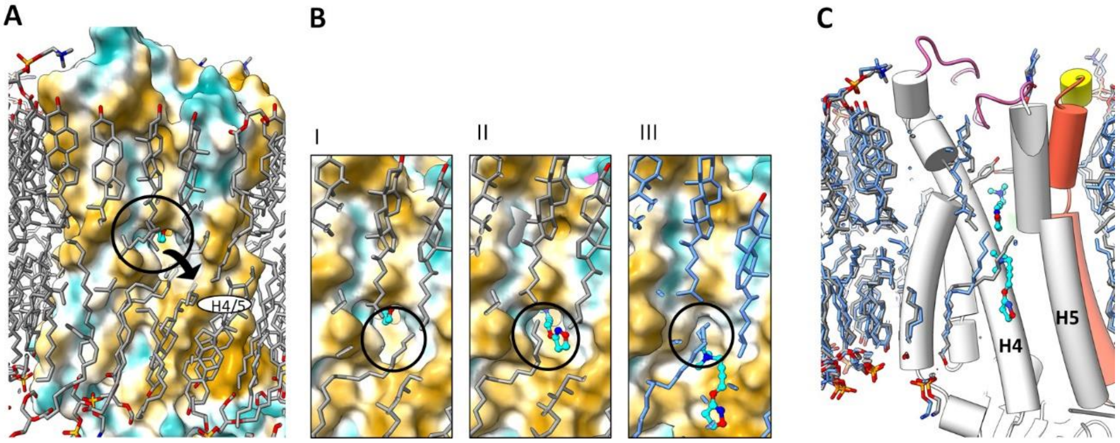


***Figure S9 Dissociation of iperoxo towards the membrane between helices 4 and 5 in a RAMD simulation in the presence of the PAM**, showing how the lipid is displaced by the ligand as it exits from the protein. (**A, B**) The protein surface colored by hydrophobicity (yellow: hydrophobic, green : hydrophilic) and the ligand shown with cyan carbon atoms; the* position from which the ligand exits is shown by the black circle and arrow; *(**A**) The protein with lipid molecules located in the vicinity of the protein, (**B**) Three dissociation steps in the RAMD trajectory: a lipid molecule covering the egress tunnel is pushed away by the ligand and then moves back after the ligand as exited; (**C**) Two frames of the same dissociation trajectory are shown with the protein displayed in cartoon representation: lipid position in the starting (protein and ligand are in a complex) and final (after ligand dissociation) frames are shown in gray and blue sticks, respectively .*
